# Effect of *neuB* deletion on biofilm formation and extracellular electron transfer in *Geobacter sulfurreducens*

**DOI:** 10.64898/2026.11.17.767340

**Authors:** Bernardo Jaramillo-Rodríguez, Eder Gómez-Escorza, Guillermo Huerta-Miranda, Alberto Hernández-Eligio, Katy Juárez

**Affiliations:** Departamento de Microbiología Molecular, Instituto de Biotecnología, Universidad Nacional Autónoma de México. Av. Universidad 2001, Col. Chamilpa, Cuernavaca, Morelos, México; Investigador por México, Secretaría de Ciencia, Humanidades, Tecnología e Innovación, Ciudad de México, México

**Keywords:** *Geobacter sulfurreducens*, biofilm, electrochemical analysis, N-acetylneuraminic acid synthesis, *neuB* gene

## Abstract

The extracellular matrix of *Geobacter sulfurreducens* biofilms is primarily composed of proteins and exopolysaccharides. The extracellular polymeric substances (EPS) matrix provides the structural scaffold necessary to facilitate extracellular electron transfer (EET); however, the specific role of its carbohydrate components remains unexplored. Sialic acids are a class of carbohydrate derivates that can modulate extracellular matrix stability in bacteria. N-acetylneuraminic acid (Neu5Ac), the most common sialic acid, is primarily synthesized by NeuB. We hypothesized that the *neuB* gene homolog in *G. sulfurreducens* (*gsu1970*), putatively involved in Neu5Ac synthesis, plays a key role in EPS organization and EET performance. Deletion of *neuB* reduced insoluble Fe(III) oxide reduction by approximately 91%. Furthermore, confocal laser scanning microscopy (CLSM) revealed that the Δ*neuB* strain formed significantly less dense EPS matrices on both hematite and fluorine-doped tin oxide (FTO) electrodes. Interestingly, bioelectrochemical analysis of FTO anodes showed that the mutant produced a 3-fold higher current density and exhibited a more negative open-circuit potential than the wild-type strain. It is important to mention that this enhanced performance on FTO is linked to a significant upregulation of *omcS*, however, more specific experiments need to be performed to establish a definitive causal relationship between *omcS* transcription levels and the observed bioelectrochemical enhancement. This transcriptional change was not observed during growth on hematite. Our results suggest that *neuB* expression contributes to maintaining the structural integrity of the WGA-reactive glycoconjugates within the EPS matrix during insoluble iron reduction. Conversely, its absence is associated with a compensatory upregulation of *omcS* on FTO electrodes. Overall, this study highlights the potential importance of putative sialic acids-related biosynthetic pathways in *Desulfuromonadia*, showing that alteration of EPS carbohydrate biosynthetic genes can markedly influence both iron reduction and the performance of electroactive biofilms behavior.

**Key points:**

- Putative *neuB* regulates EPS matrix integrity in *G. sulfurreducens*
- Loss of *neuB* severely cuts insoluble Fe(III) oxide reduction by 91%
- Δ*neuB* mutant shows a 3× higher current density on FTO anodes via *omcS* upregulation

## Introduction

Sialic acids are a family of nine-carbon monosaccharides belonging to the group of nonulosonic acids (NulOs) (Ghosh 2020). Typically located at the terminal positions of glycans and other glycoconjugates, these sugars confer critical functional properties on the extracellular matrix, including protection against enzymatic degradation and the modulation of cell-cell and cell-surface interactions. Consequently, sialic acids have been implicated in biofilm matrix organization and cell–surface interactions, and are associated with changes in biofilm structural properties of microbial biofilms (Ghosh 2020; Kleikamp et al. 2020; Paleček et al. 2015; Cardona et al. 2025).

Although the synthesis and functions of sialic acids have been extensively documented in pathogenic bacteria, where they contribute to molecular mimicry, host immune evasion, and biofilm-associated virulence, their role in environmental microorganisms remains largely unexplored (Almagro-Moreno and Boyd 2009). However, increasing evidence indicates that sialic acids and related nonulosonic acids are also present in biofilms produced by environmental microorganisms (Pinel et al. 2020; Boleij et al. 2020; Vieira et al. 2021; Chen et al. 2023). These findings suggest that sialic acids and related nonulosonic acids play functional roles in extracellular matrix modification and structural organization in environmental biofilms.

*Geobacter sulfurreducens* is a Gram-negative, anaerobic bacterium belonging to the class *Desulfuromonadia* (Lovley et al. 2011). Because *G. sulfurreducens* can use insoluble electron acceptors, such as Fe(III) and Mn(IV) oxides, as well as artificial electrodes in bioelectrochemical systems through extracellular electron transfer (EET), it has become a model organism for studies of bioremediation and bioelectricity generation. The formation of electroactive biofilms is fundamental to this EET process (Bond and Lovley 2003; Xu et al. 2025). The development and electrochemical performance of these biofilms depend on a coordinated matrix composed of conductive pili, c-type cytochromes, and exopolysaccharides (Lovley et al. 2011; Reguera and Kashefi 2019). Within this matrix, EPS not only promotes cell-to-cell adhesion and surface attachment but also anchors essential electroactive proteins, including outer-membrane cytochromes, thereby positioning them for efficient electron transfer (Rollefson et al. 2011; Zhuang et al. 2020, 2022).

Genomic analysis of *G. sulfurreducens* has identified a cluster of approximately 35 genes putatively involved in the synthesis, transport, and modification of EPS matrix components (Jaramillo-Rodríguez et al. 2023). This *eps* cluster includes genes predicted to participate in N-acetylneuraminic acid (Neu5Ac) biosynthesis (*gsu1970* [*neuB*], *gsu1972* [*neuA*], and *gsu1973* [*neuD*]) (Ghosh 2020; Jaramillo-Rodríguez et al. 2023). Previous studies showed that deleting *gsu1771*—which encodes a transcriptional repressor of this *eps* cluster— results in the upregulation of the entire cluster, altering biofilm phenotypes and enhancing extracellular electron transfer (EET) capacity (Hernández-Eligio et al. 2022; Jaramillo-Rodríguez et al. 2023; Rodríguez-Torres et al. 2025). However, because GSU1771 controls a large multi-gene cluster, the specific contribution of individual biosynthetic pathways within this cluster remains unknown.

To dissect the functional role of sialic acids-related pathways within this cluster without disrupting the entire EPS machinery, we targeted *gsu1970* (predicted *neuB*). In bacteria such as *E. coli*, the NeuB enzyme catalyzes the condensation of D-ManNAc with phosphoenolpyruvate, representing the predicted committed step in *de novo* Neu5Ac backbone synthesis. Therefore, deleting *neuB* provides a targeted genetic strategy to test the hypothesis of whether disrupting this predicted biosynthetic step impacts the structural organization of EPS matrix and EET performance in *G. sulfurreducens*.

## Materials and methods

### Bacterial strains and culture conditions

The *G. sulfurreducens* strains used in this study are listed in Supplementary Table 1. *G. sulfurreducens* strains were routinely cultured in NBAF medium (using acetate and fumarate as the electron donor and electron acceptor, respectively), as previously described (Coppi et al. 2001). To evaluate Fe(III) reduction, *G. sulfurreducens* strains were cultivated in either ferric citrate or Iron-Gel medium (Coppi et al. 2001). The ferric citrate medium contained (per liter): 13.7 g ferric citrate, 2.5 g NaHCO_3_, 0.25 g NH_4_Cl, 0.6 g NaH_2_PO_4_·H_2_O, 0.1 g KCl, 1.0 mL Na_2_SeO_4_ (100 mM), 10 mL DL vitamin mix (100×), and 10 mL NB mineral mix (100×), supplemented with 20 mM sodium acetate (Coppi et al. 2021). Fe(III) oxide gel was prepared by dissolving 108 g of FeCl_3_ in 1 L of deionized water under constant magnetic stirring. The pH of the solution was adjusted to 7.0 using 10 N NaOH, stirred for 30 min, and readjusted if necessary. The resulting precipitate was harvested by centrifugation at 3,500×g for 20 min, washed by resuspension in Milli-Q water, and centrifuged three times to remove residual salts. To determine the total Fe concentration of each batch, 100 μL of the purified Fe-gel was digested with 4.7 mL of 0.5 N HCl and 200 μL of 6 N hydroxylamine monohydrochloride, incubated overnight at 4 °C in the dark, and total Fe was quantified using the ferrozine assay method (Lovley and Phillips 1988). Finally, Iron-gel medium was prepared containing (per liter): 100 mM synthesized Fe(III) oxide gel, 0.06 g NaH_2_PO_4_, 2.5 g NaHCO_3_, 0.1 g KCl, 0.25 g NH_4_Cl, 10 mL DL vitamin mix (100×), and 10 mL NB mineral mix (100×), supplemented with 20 mM sodium acetate. Anoxic conditions were established by purging the media with an N_2_/CO_2_ gas mixture (80:20, v/v) to remove residual O_2_. Cultures were incubated statically at either 30 °C or 25 °C. When required, the media were supplemented with kanamycin (Km, 300 µg/mL) or spectinomycin (Sp, 70 µg/mL) for selective growth.

To examine the physical and electrochemical properties of *G. sulfurreducens* biofilms, cells were acclimated for surface attachment on fluorine-doped tin oxide (FTO) and hematite-coated FTO (Fe_2_O_3_-FTO) electrodes at 25 °C through a two-week serial transfer procedure in fresh fumarate-containing NBAF medium (Hernández-Eligio et al. 2022). Briefly, cultures were incubated statically with submerged electrodes for one week to allow initial biofilm colonization. At the end of each week, the culture bottles were agitated to detach the attached biofilm biomass, resuspending both biofilm-derived and planktonic cells. This mixed cell suspension was then used to inoculate fresh NBAF medium containing a clean electrode support. After two consecutive weekly acclimation cycles, the resuspended cells were inoculated into fresh NBAF medium with new electrode supports to allow the growth of experimental biofilms for exactly 48 h or 72 h at 25 °C prior to open-circuit or electrochemical analysis. It is important to note that under these conditions, fumarate serves as the primary soluble electron acceptor to support biofilm colonization prior to open-circuit or electrochemical analysis. Mature biofilms grown on the electrodes at 25 °C for 48 or 72 h were subsequently analyzed.

*Escherichia coli* was cultured in LB medium at 37 °C with orbital shaking (200 rpm) and supplemented with Km (30 µg/mL) or Sp (35 µg/mL), as appropriate. *E. coli* strains S17-1 and MC1061 were used for conjugation with *G. sulfurreducens* and for plasmid transformation, maintenance, and isolation, respectively (Supplementary Table 1).

### DNA extraction and manipulation

Genomic DNA, plasmids, and PCR products were purified using the DNeasy Blood and Tissue Kit (Qiagen, Germany), the High Pure Plasmid Isolation Kit (Roche, Germany), and the GeneJET PCR Purification Kit (Thermo Fisher Scientific, Vilnius, Lithuania), respectively. *E. coli* transformations and all other routine DNA manipulations were performed according to standard molecular biology protocols (Sambrook et al. 1989).

### Construction of the Δ*neuB* mutant strain

The Δ*neuB* mutant strain was generated by scarless mutagenesis using the suicide plasmid pK18mobSacB, as described by Chan et al. (2015). To construct the mutant allele, the regions flanking *neuB* were amplified by PCR from *G. sulfurreducens* DL1 genomic DNA using 2× Phusion Flash High-Fidelity PCR Master Mix (Thermo Fisher Scientific, Vilnius, Lithuania). A 983-bp upstream fragment was amplified using primers Primer1-HindIII-neuB and Primer2-neuB, whereas a 1,510-bp downstream fragment was amplified using Primer3-neuB and Primer4-EcoRI-neuB (Supplementary Table 1). The purified amplicons were subsequently used as megaprimers in an overlap-extension PCR strategy to generate a 2,453-bp Δ*neuB* chimeric fragment.

The resulting chimeric PCR product was digested with *Eco*RI and *Hin*dIII and ligated into pK18mobsacB (Supplementary Table 1), which had been digested with the same restriction enzymes, generating pK18mobsacB-ΔneuB. This plasmid was introduced into *E. coli* S17-1 by electroporation and subsequently transferred to the recipient *G. sulfurreducens* DL1 strain by conjugation. Briefly, 1 mL of a late-exponential-phase NBAF culture of *G. sulfurreducens* was pelleted together with 1 mL of an *E. coli* S17-1/pK18mobsacB-ΔneuB culture. The bacterial mixture was deposited onto 0.22-µm-pore-size filters placed on acetate-fumarate agar plates inside an anaerobic chamber and incubated for 4 h. Cells were then streaked onto acetate-fumarate agar plates containing kanamycin to select single-crossover recombinants. Double-crossover scarless deletion mutants were subsequently counterselected on acetate-fumarate anaerobic plates supplemented with 10% (w/v) sucrose. Candidate mutants were confirmed by PCR using primers Primer1-HindIII-neuB and Primer4-EcoRI-neuB together with Taq DNA Polymerase (Thermo Fisher Scientific, Vilnius, Lithuania) (Supplementary Fig. 1).

### EPS extraction

EPS extraction was performed according to the protocol described by Cao et al. (2011) with minor modifications. *G. sulfurreducens* DL1 and Δ*neuB* biofilms were grown statically on glass surfaces in NBAF medium at 25 °C for 72 h. Biofilms were harvested, resuspended in a 1:1 (v/v) mixture of 0.9% NaCl and 2% EDTA, and incubated on ice for 3 h with gentle agitation. The suspension was then centrifuged at 5,000 × g for 20 min at 4 °C, and the supernatant containing the extracted EPS fraction was collected for further analysis. Total protein content in the extracted EPS fraction was quantified using the Bradford assay (Bio-Rad, USA) with bovine serum albumin (BSA) as the standard to evaluate matrix yield across strains.

### Spectroscopic detection of *c*-type cytochromes in the extracellular matrix

To evaluate the presence and redox state of *c*-type cytochromes within the extracted EPS fraction, UV-Vis absorption spectra were recorded using a Beckman Coulter spectrophotometer (California, USA). Extracts were first analyzed under oxidizing conditions (exposure to ambient oxygen) to monitor the Soret (γ) peak at approximately 408 nm, characteristic of oxidized cytochrome *c* (Nakamura et al. 2009; Zhuang et al. 2022). Samples were subsequently reduced under anoxic conditions by adding 100 mM sodium dithionite prepared in 100 mM HEPES buffer (Jones et al. 1983). The spectra of the reduced samples were recorded to detect the characteristic shift of the Soret band (∼419 nm) and the appearance of the characteristic cytochrome *c* α (∼555 nm) and β (∼525 nm) peaks. To reliably compare the relative cytochrome-associated spectral signal across strains, absorbance data were normalized using the OD600 values of the biofilm cells (measured from a 1:10 dilution of a cell suspension taken prior to EPS extraction) and cross-referenced with total EPS protein concentrations.

### Analytical techniques

Fe(II) generated during soluble and insoluble Fe(III) reduction was quantified using the standard ferrozine spectrophotometric assay (Lovley and Phillips, 1988).

### Electrochemical characterization of biofilms

Electrochemical analyses were performed in a custom-built three-electrode electrochemical cell consisting of an Ag/AgCl reference electrode (saturated KCl), a platinum wire counter electrode, and biofilm-coated FTO electrodes as the working electrode (Supplementary Fig. 2). The electrodes were immersed in an anaerobic basal medium containing 30 mM NaHCO_3_, 5 mM NH_4_Cl, and 50 µM NaH_2_PO_4_·H_2_O under continuous N_2_ sparging, either in the absence or presence of 20 mM sodium acetate as the electron donor. Electrochemical measurements were performed using a Metrohm DropSens potentiostat (Asturias, Spain) controlled with DropView 8400 software. Open-circuit potential (OCP) was recorded every 0.5 s for 900 s. Cyclic voltammetry (CV) was performed at a scan rate of 0.001 V/s over a potential range of −0.8 to +0.5 V (vs. SHE). Square-wave voltammetry (SWV) was conducted using a step potential of 0.001 V, a modulation amplitude of 0.01 V, and a frequency of 37 Hz over the same potential range to resolve individual redox peaks. All potentials were converted to and reported on the standard hydrogen electrode (SHE) scale (SHE = Ag/AgCl + 0.199 V). The current density was estimated by dividing the current by the electroactive area of each electrode. The electroactive area of the support electrodes was calculated by the chronoamperometry method and the Cottrell equation (Cummings et al. 1998).

To describe the kinetic behavior of electroactive biofilms, non-turnover and turnover cyclic voltammograms were modeled using a modified Nernst–Monod equation based on a single-step, one-electron transfer process coupled to substrate utilization (Marcus et al. 2007; Torres et al. 2008, 2010):

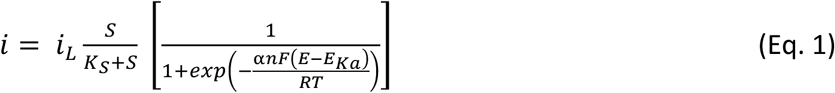

The Nernst-Monod equation (Eq. 1) is a hybrid of the Monod and Butler-Volmer equations, parameterized for bioelectrochemical systems: where i is the steady-state current, i_L_ is the maximum current, S is the substrate concentration (0.011 mol/L, measured in terms of COD at 48 h in the WT NBAF medium; Dhar et al. 2016), K_S_ is the half-saturation constant (1.2 × 10^−4^ mol/L), α is the charge transfer coefficient, E is the electrode potential (V vs. SHE), E_Ka_ is the half-wave potential (≍ E^0^’, V vs. SHE) representing the predominant outer-membrane *c-type* cytochrome redox network active at the biofilm–anode interface, R is the gas constant (8.314 Jmol^−1^K^−1^), T is the absolute temperature (298.15 K), n is the number of electrons, and F is the Faraday constant (96,485 Cmol^−1^). Prior to parameter estimation, non-faradaic background capacitive currents were subtracted by averaging forward and backward scans under non-turnover conditions. Ohmic drop compensation was applied using uncompensated solution resistance values obtained from electrochemical impedance spectroscopy. Non-linear least-squares fitting was performed within the potential range of −0.50 to +0.50 V (vs. SHE). Goodness-of-fit was evaluated using the residual sum of squares (RSS) and the coefficient of determination (R^2^>0.98) (Cholula-Calixto et al. 2026).

### Biofilm analysis by confocal laser scanning microscopy

Biofilm structural characteristics and cell viability were examined using confocal laser scanning microscopy (CLSM). For viability assays, *G. sulfurreducens* DL1 and Δ*neuB* biofilms were grown on FTO electrodes for 48 h at 25 °C and stained using the LIVE/DEAD® BacLight Bacterial Viability Kit (Invitrogen, Thermo Fisher Scientific, Eugene, Oregon, USA), as previously described (Huerta-Miranda et al. 2019). Electrodes were gently removed from the culture tubes, washed with an anaerobic solution containing 0.9% NaCl and 100 mM cysteine (1:1, v/v), and incubated for 10 min with a mixture of SYTO9 (3.34 mM) and propidium iodide (20 mM).

To analyze extracellular matrix components, biofilms were cultured for 72 h on either uncoated FTO or Fe_2_O_3_-FTO electrodes (Rodríguez-Torres et al. 2025). Extracellular proteins were stained with SYPRO® Orange (5×) (Invitrogen, Thermo Fisher Scientific, Eugene, Oregon, USA). Extracellular glycoconjugates were stained with Wheat Germ Agglutinin-Alexa Fluor 633 (WGA-AF633; 50 µg/mL; Invitrogen, Thermo Fisher Scientific, Eugene, Oregon, USA) which binds to both N-acetylglucosamine (GlcNAc) and N-acetylneuraminic acid (Neu5Ac) residues, to monitor WGA-reactive glycoconjugates within the matrix, and with Concanavalin A-Alexa Fluor 633 (ConA-AF633; 100 µg/mL; Invitrogen, Thermo Fisher Scientific, Eugene, Oregon, USA) to detect α-D-glucose- and α-D-mannose-rich glycoconjugates.

CLSM imaging was performed using an Olympus FV1000 microscope with a LUMFLN 60×/1.10 W water-immersion objective. To allow reliable quantitative fluorescence intensity comparisons, all imaging parameters were kept strictly identical across all experimental groups and control samples. Fluorescent signals were acquired in sequential XYZ scanning mode (unidirectional scan, scan speed 4.0). The pinhole diameter was set to 250 µm. Laser power/irradiance lines were set to 488 nm (0.00219 mW), and 559 nm (0.221 mW). Channel 1 (SYTO9) were excited at 488 nm and detected at 520 nm (PMT voltage: 675 V, detector offset: 10). Channel 2 (SYPRO Orange) were excited at 559 nm using and detected at 618 nm (PMT voltage: 825 V, detector offset: 10). Channel 3 (WGA-AF633, ConA-AF633) were excited at 559 nm and detected at 647 nm (PMT voltage: 800 V, detector offset: 10). Images were captured at 800 × 800 pixel resolution (12-bit depth, pixel size of 0.265 µm/pixel, 211.74 µm field of view) using Line-Kalman filtering (count = 4) for noise reduction. Z-stacks spanning the entire biofilm thickness were collected for quantitative analysis. Images were processed and quantitatively analyzed using Fiji (version 2.9.0) (Schindelin et al. 2012).

### Complementation of the Δ*neuB* mutant strain

For genetic complementation, the expression plasmid pRneuB was constructed by placing the wild-type *neuB* gene under the control of the constitutive *G. sulfurreducens recA* promoter. The *recA* regulatory region (RRrecA), including its ribosome-binding site (RBS), was amplified by PCR using primers EcoRI-recAPromFw and NdeI-recAPromRv and cloned into pJET1.2 (CloneJET PCR Cloning Kit, Thermo Fisher Scientific, Vilnius, Lithuania) to generate pRRrecA (Supplementary Table 1). The *neuB* coding sequence was amplified using primers NdeI-neuBFw and HindIII-neuBRv. The *neuB* amplicon was digested with *Nde*I and *Hind*III and ligated into the corresponding sites of pRRrecA, generating pRRrecA-neuB. The RRrecA-neuB expression cassette (1,224 bp) was excised with *Eco*RI and *Hind*III and inserted into the broad-host-range vector pRG5.1 (Kim et al. 2005) to generate the complementation plasmid pRneuB (Supplementary Table 1). After verification by restriction analysis and sequencing, pRneuB was introduced into the Δ*neuB* mutant by electroporation. Transformants were selected on spectinomycin-containing anaerobic plates, and plasmid maintenance was confirmed by restriction digestion.

### RNA extraction and purification

To determine transcriptional profiles, *G. sulfurreducens* DL1 and Δ*neuB* strains were grown anaerobically in NBAF medium at 25 °C. Biofilms were harvested from FTO electrodes after 48 h of growth or from Fe_2_O_3_-FTO electrodes after 72 h. Electrodes were removed, and the attached biomass was scraped into 2 mL of ice-cold phosphate buffer. To preserve transcript stability, 100 µL of RNAlater (Thermo Fisher Scientific, Vilnius, Lithuania) was added immediately before cell pelleting (3,000 rpm for 2 min). Total RNA was extracted using the RNeasy Mini Kit (Qiagen, Germany) according to the manufacturer’s instructions. Residual genomic DNA was removed by DNase I (Thermo Fisher Scientific, Vilnius, Lithuania) digestion. RNA concentration and purity were determined spectrophotometrically (NanoDrop, Thermo Scientific, USA), and RNA integrity was assessed by electrophoresis on a 1% agarose gel prepared in 0.5× TBE buffer.

### Quantitative reverse transcription PCR

Complementary DNA (cDNA) was synthesized from 1 µg of purified total RNA using the RevertAid First Strand cDNA Synthesis Kit (Thermo Fisher Scientific, Vilnius, Lithuania) according to the manufacturer’s instructions. Reverse transcription was primed with the gene-specific reverse primers listed in Supplementary Table 1. Quantitative reverse transcription PCR (RT-qPCR) was performed using a Rotor-Gene 2plex HRM instrument (Qiagen, Hilden, Germany) and Maxima SYBR Green/ROX qPCR Master Mix (Thermo Fisher Scientific, Vilnius, Lithuania). The expression levels of the outer-membrane cytochrome genes *omcB, omcC, omcE, omcS, omcZ, omcM,* and *omcF*, the periplasmic cytochrome gene *pgcA*, and the structural pilin gene *pilA* were evaluated. Relative gene expression was calculated using the 2^-ΔΔCT^ method, with *recA* serving as the reference gene. All reactions were performed in technical triplicate using duplicate biological samples from independent experiments.

## Results

### The *neuB* gene is part of the putative Neu5Ac biosynthesis pathway in *G. sulfurreducens*

Comparative genomic analysis identified a putative Neu5Ac biosynthesis pathway in *G. sulfurreducens* that shares structural similarities with those described in *E. coli* and *Neisseria meningitidis* (Supplementary Fig. 3a). In *E. coli*, Neu5Ac biosynthesis begins with UDP-α-D-GlcNAc, which is converted to D-ManNAc by the N-acetylglucosamine 2-epimerase NeuC (SiaA in *N. meningitidis*). NeuB (SiaC in *N. meningitidis*) subsequently catalyzes the condensation of D-ManNAc with phosphoenolpyruvate to produce Neu5Ac (Petersen et al. 2000; Vann et al. 2004; Ghosh 2020).

In the *G. sulfurreducens* genome, *gsu2243* was identified as a homolog of *neuC/siaA*, whereas a homolog of *neuB* (*gsu1970*) is located within a distinct genomic region comprising approximately 35 genes (Supplementary Fig. 3b). This region also contains *asnB* (*gsu1953*), which encodes a putative asparagine synthetase, and *gsu1989*, encoding a sigma-54-dependent response regulator (Methé et al. 2003). Throughout this study, this genomic region is referred to as the *eps* cluster. The *eps* cluster contains multiple open reading frames predicted to participate in the synthesis, export, and surface modification of EPS (Jaramillo-Rodríguez et al. 2023). Among these genes are *gsu1972*, a homolog of *neuA* encoding a putative CMP-sialic acid synthetase, and *gsu1973*, a homolog of *neuD* encoding a putative O-acetyltransferase involved in Neu5Ac modification (Ghosh 2020).

### Effect of the *neuB* mutation on *G. sulfurreducens* biofilms

The predicted NeuB protein of *G. sulfurreducens* shares 31.86% and 32.84% amino acid sequence identity with the NeuB homologs from *N. meningitidis* and *E. coli*, respectively. Multiple sequence alignment showed that the *G. sulfurreducens* enzyme retains 15 of the 16 highly conserved catalytic residues required for NeuB activity in these organisms (Supplementary Fig. 4a) (Gunawan et al. 2005). An in silico structural analysis of the *G. sulfurreducens* NeuB protein using AlphaFold predicts an N-terminal (β/α)_8_-barrel (TIM barrel) and a C-terminal domain structurally resembling type III ice-binding antifreeze proteins, consistent with its homologs in *N. meningitidis* and *E. coli* (Gunawan et al. 2005). Furthermore, NeuB is predicted to harbor a Mn^2+^-binding motif essential for its enzymatic activity (Supplementary Fig. 4b). To examine the role of *neuB* in Neu5Ac production, a scarless Δ*neuB* mutant was generated (Supplementary Fig. 1). Biofilms of the wild-type DL1 strain and the Δ*neuB* mutant were grown on FTO electrodes for 48 h and stained with WGA-AF633 to visualize Neu5Ac-containing glycoconjugates (Fig. 1). Cell viability was simultaneously evaluated using the LIVE/DEAD® BacLight staining kit (Fig. 2). CLSM analysis revealed a marked reduction in WGA-AF633 fluorescence throughout the extracellular matrix of the Δ*neuB* biofilm relative to the wild-type strain, indicating a decrease in WGA-reactive glycoconjugates (containing Neu5Ac/GlcNAc residues) (Fig. 1a). Quantitative image analysis using Fiji demonstrated an approximately 3-fold decrease in fluorescence intensity in the mutant biofilms (Fig. 1b). Introduction of the complementation plasmid pRneuB restored extracellular Neu5Ac-reactive fluorescence to levels comparable to those of the wild type, whereas the empty vector pRG5.1 did not restore the phenotype (Fig. 1a,c). Cell viability also differed between strains. Wild-type biofilms contained 90.22 ± 2.49% viable cells, whereas viability decreased to 77.92 ± 0.74% in the Δ*neuB* biofilms (Fig. 2).

**Fig. 1.**
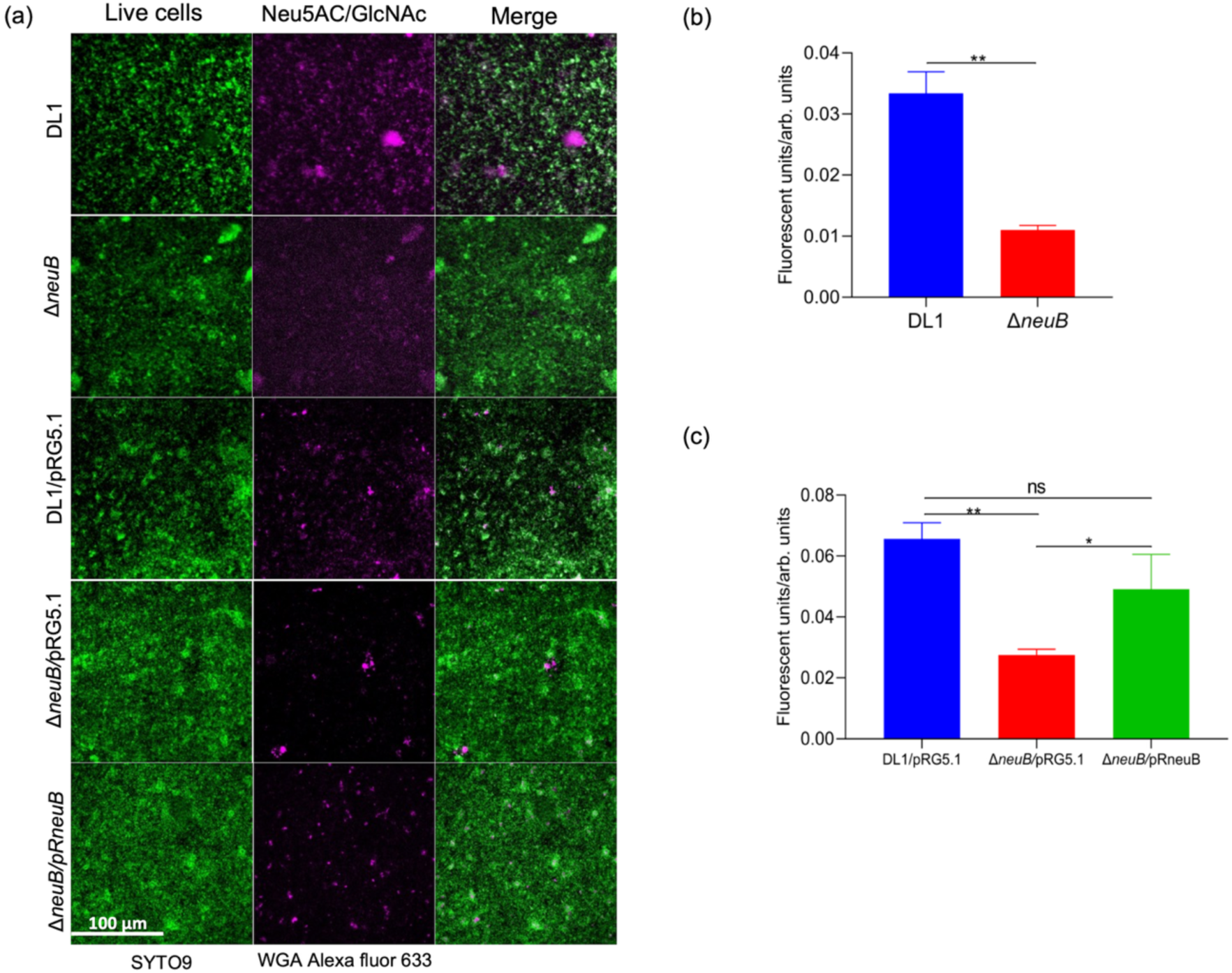
Detection of Neu5Ac/GlcNAc residues in *G. sulfurreducens* biofilms by CLSM. **(a)** Representative CLSM images of biofilms stained with SYTO9 (green, viable cells) and WGA-AF633 (magenta, extracellular matrix glycans containing Neu5Ac/GlcNAc residues). Individual fluorescence channels (viable cells and Neu5Ac/GlcNAc) and the merged images are shown for the DL1, Δ*neuB*, DL1/pRG5.1 (empty vector), Δ*neuB*/pRG5.1 (empty vector), and Δ*neuB*/pR*neuB* (complementation vector carrying the wild-type *neuB* gene) strains. Scale bar = 100 µm. **(b)** Quantification of WGA-AF633 fluorescence in DL1 and Δ*neuB* biofilms. **(c)** Quantification of WGA-AF633 fluorescence in the empty vector controls and the complemented strain. In **(b)** and **(c)**, WGA-AF633 fluorescence is expressed in arbitrary units for a biofilm volume of 4 × 10^5^ μm^3^ and normalized to SYTO9 fluorescence. Bars represent the mean of two independent experiments, and error bars indicate the variability between replicates. Statistical significance was determined by one-way ANOVA followed by Šídák’s multiple-comparisons test (ns, *P* > 0.05; * *P* < 0.05; ** *P* < 0.01)

**Fig. 2.**
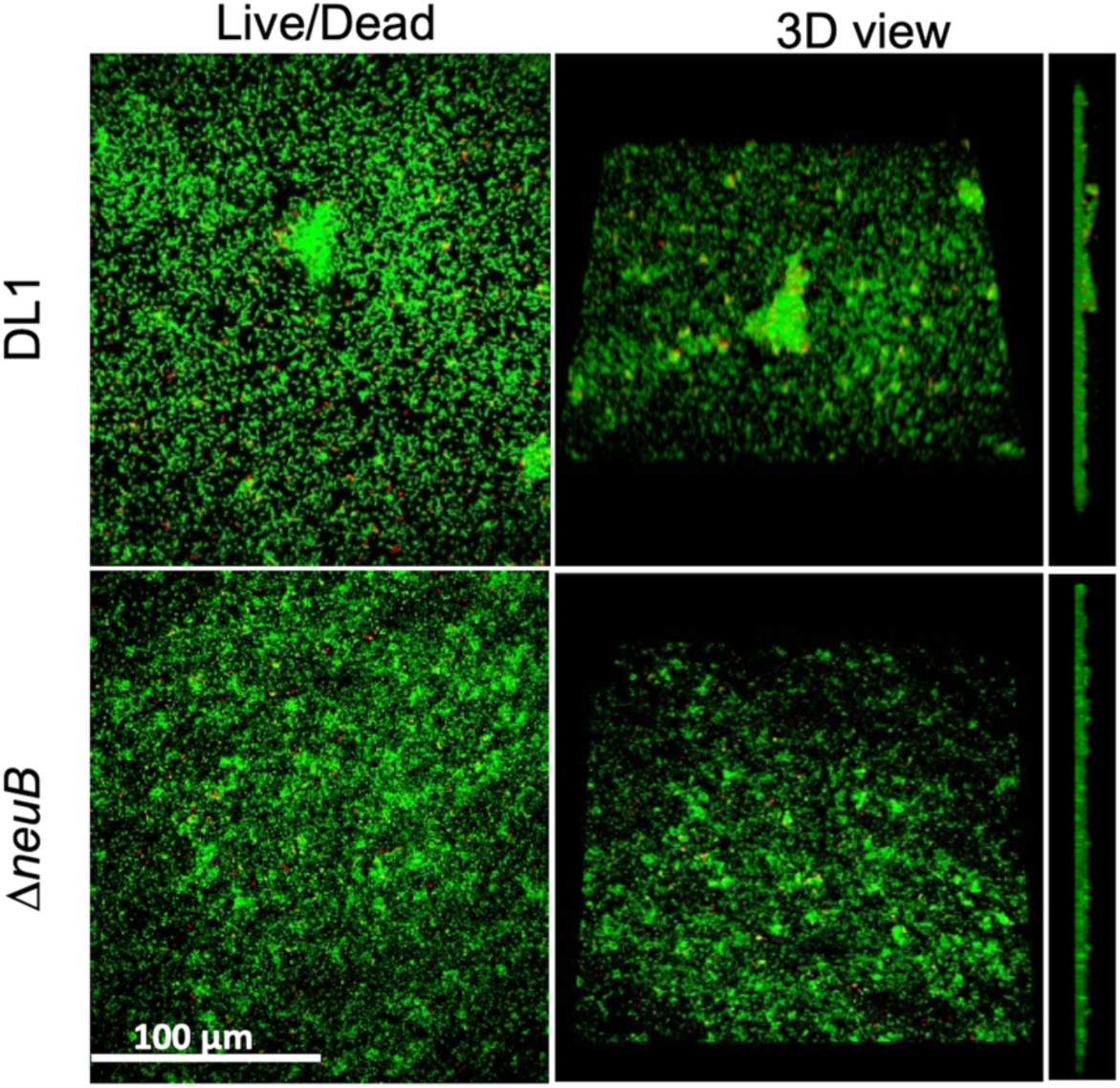
Analysis of cell viability in DL1 and Δ*neuB* biofilms by CLSM. Representative CLSM images of biofilms stained with the LIVE/DEAD® BacLight Bacterial Viability Kit are shown as top-view projections together with three-dimensional reconstructions generated using Fiji, including oblique and lateral views. Green fluorescence indicates viable cells, whereas red fluorescence indicates dead cells. Scale bar = 100 µm

### Effect of the Δ*neuB* mutation on planktonic growth and Fe(III) reduction

Deletion of *neuB* did not affect growth when fumarate or soluble Fe(III)-citrate served as the terminal electron acceptor, as both the wild-type and mutant strains displayed comparable growth kinetics (Fig. 3a,b). In contrast, substantial differences were observed during growth with insoluble Fe(III) oxide. Under these conditions, the Δ*neuB* mutant exhibited markedly impaired Fe(III) reduction (Fig. 3c). At the end of the incubation period, the wild-type strain produced 25.14 mM Fe(II), whereas the Δ*neuB* mutat generated only 2.26 mM Fe(II), representing an approximately 91% reduction in insoluble Fe(III) reduction capacity (Fig. 3c).

**Fig. 3.**
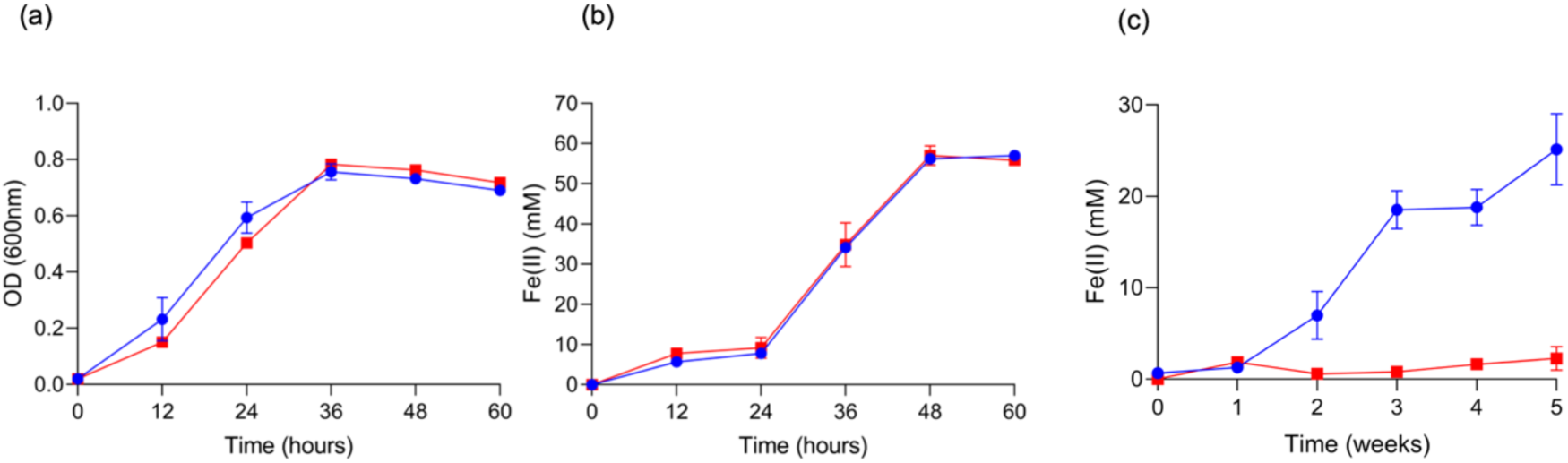
Growth and Fe(III) reduction by DL1 and Δ*neuB* strains. **(a)** Growth in NBAF medium with acetate and fumarate as the electron donor and acceptor, respectively. **(b)** Reduction of soluble Fe(III). **(c)** Reduction of insoluble Fe(III). In **(a)**, **(b)**, and **(c)**, the blue line represents the DL1 wild-type strain, and the red line represents the Δ*neuB* mutant strain. Data points represent the mean of three independent experiments, and error bars indicate standard deviation

To determine whether deletion of *neuB* altered extracellular cytochrome accumulation, total EPS fractions were extracted and analyzed by UV-Vis spectroscopy under oxidized and sodium dithionite-reduced conditions. Both strains exhibited nearly identical absorbance spectra, including the characteristic cytochrome *c* peaks observed after reduction (Supplementary Fig. 5), indicating that extracellular cytochrome accumulation was not detectably altered by the mutation.

### Effect of *neuB* deletion on the expression of genes involved in Fe(III) reduction

To examine transcriptional changes during biofilm development on mineral surfaces, expression levels of key EET genes were evaluated in biofilms grown on hematite-coated surfaces (Fe_2_O_3_-FTO) in NBAF (acetate-fumarate) medium under open-circuit potential (OCP). Because fumarate was present in the medium to support baseline colonization and growth prior to and during OCP measurements, these transcriptional profiles represent the physiological state of biofilms developed on hematite surfaces under OCP rather than a strict, sole-source insoluble iron-respiring state. Under these conditions, RT-qPCR profiles revealed that expression levels of most core EET components remained closely comparable to the wild-type control (Huerta-Miranda et al. 2023) (Fig. 4). The expression levels of the outer-membrane cytochrome genes *omcB, omcC, omcE, omcS, omcZ, omcM,* and *omcF*, together with the extracellular cytochrome gene *pgcA* and the pilin gene *pilA*, were quantified by RT-qPCR (Mehta et al. 2005; Reguera et al. 2005, 2007; Holmes et al. 2006; Inoue et al. 2010; Lovley et al. 2011; Qian et al. 2011; Liu et al. 2014; Zacharoff et al. 2017; Jiang et al. 2023). The RT-qPCR profiles revealed that most core components of the EET machinery remained transcriptionally stable, with expression levels closely mimicking those of the wild type control (Fig. 4). Overall, the transcriptional profiles of the Δ*neuB* mutant closely resembled those of the wild-type strain (Fig. 4). Only minor changes were detected, including a slight decrease in *omcS* expression (relative expression ≈ 0.86) and modest increases in *pgcA* (≈1.13) and *omcM* (≈1.37). Expression of *pilA* remained essentially unchanged (relative expression ≈0.94). Because fumarate supported baseline respiratory growth during biofilm development, the core EET transcriptional machinery remained largely unaffected (Fig. 4), highlighting that the severe phenotype observed in strict Fe(III)-only media (Fig. 3c) stems from structural EPS defects rather than transcriptional silencing of cytochromes.

**Fig. 4.**
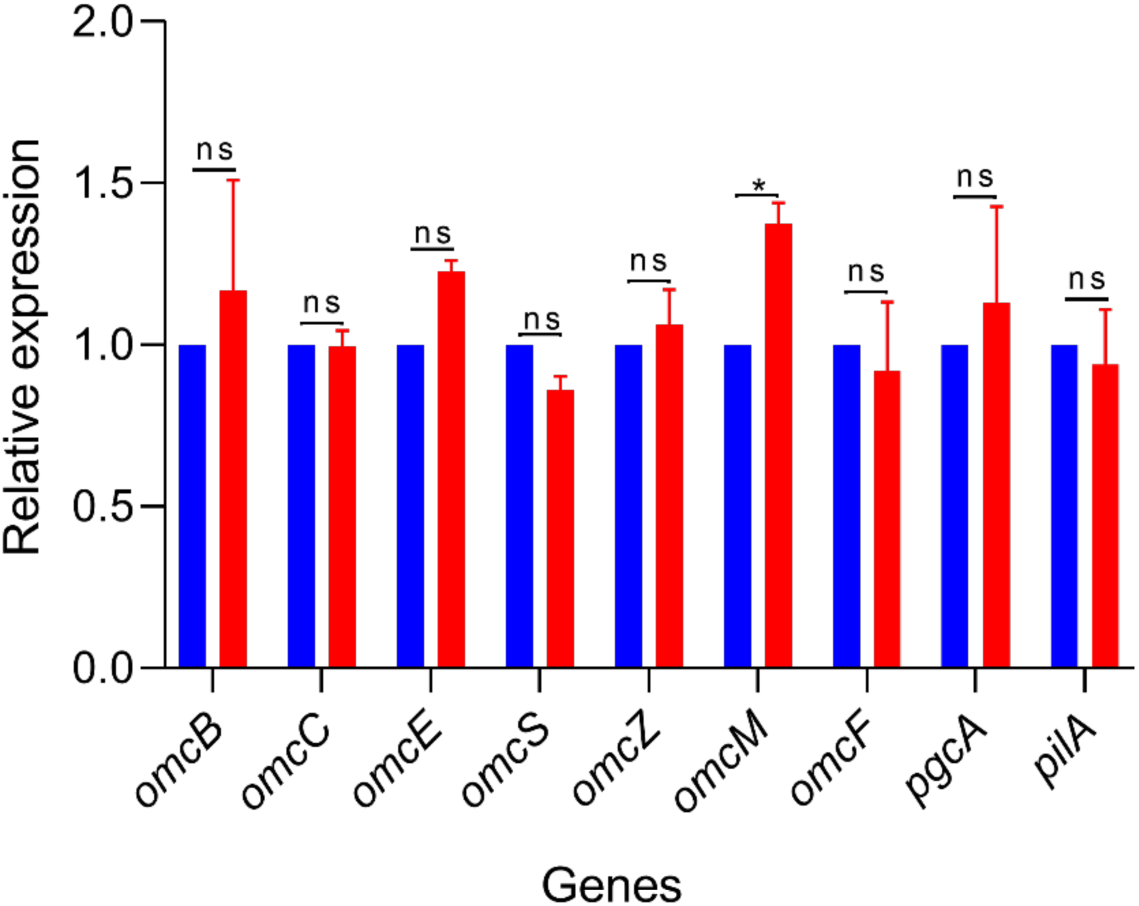
Analysis of gene expression involved in Fe(III) reduction in biofilms grown on hematite (Fe_2_O_3_-FTO). Analysis of gene expression in *G. sulfurreducens* DL1 and Δ*neuB* biofilms acclimated by serial transfer for 2 weeks and grown on hematite-coated FTO (Fe_2_O_3_-FTO) electrodes in acetate-famarate (NBAF) medium under open-circuit potential (OCP) conditions. The relative expression levels of genes encoding the cytochromes OmcB, OmcC, OmcE, OmcS, OmcZ, OmcM, OmcF, and PgcA, as well as the pilin subunit PilA, are shown. Blue bars represent gene expression levels in the wild-type DL1 strain, whereas red bars represent expression levels in the Δ*neuB* mutant strain. Error bars indicate the standard deviation of two independent experiments, and each experiment has technical triplicates per gene. Statistical significance was assessed using independent unpaired *t*-tests for each comparison without correction for multiple comparisons (ns, *P* > 0.05; * *P* < 0.05)

### Effect of the Δ*neuB* mutation on biofilm architecture during Fe(III) reduction

To examine the structural organization of biofilms during Fe(III) reduction, biofilms formed on Fe_2_O_3_-FTO electrodes were analyzed by CLSM following staining of extracellular proteins, extracellular glycoconjugates, and viable cells (Fig. 5). Wild-type biofilms formed continuous, confluent structures that uniformly covered the Fe_2_O_3_-FTO surface (Fig. 5a). The Δ*neuB* mutant developed biofilms of comparable thickness (12.1 μm) to those of the wild-type strain (12.5 μm). Despite similar thicknesses, quantitative fluorescence analysis revealed significant differences in matrix composition. The Δ*neuB* biofilms exhibited reduced extracellular protein staining and a substantially greater decrease in EPS-associated fluorescence relative to the wild-type strain (Fig. 5b), indicating a marked reduction in extracellular matrix abundance.

**Fig. 5.**
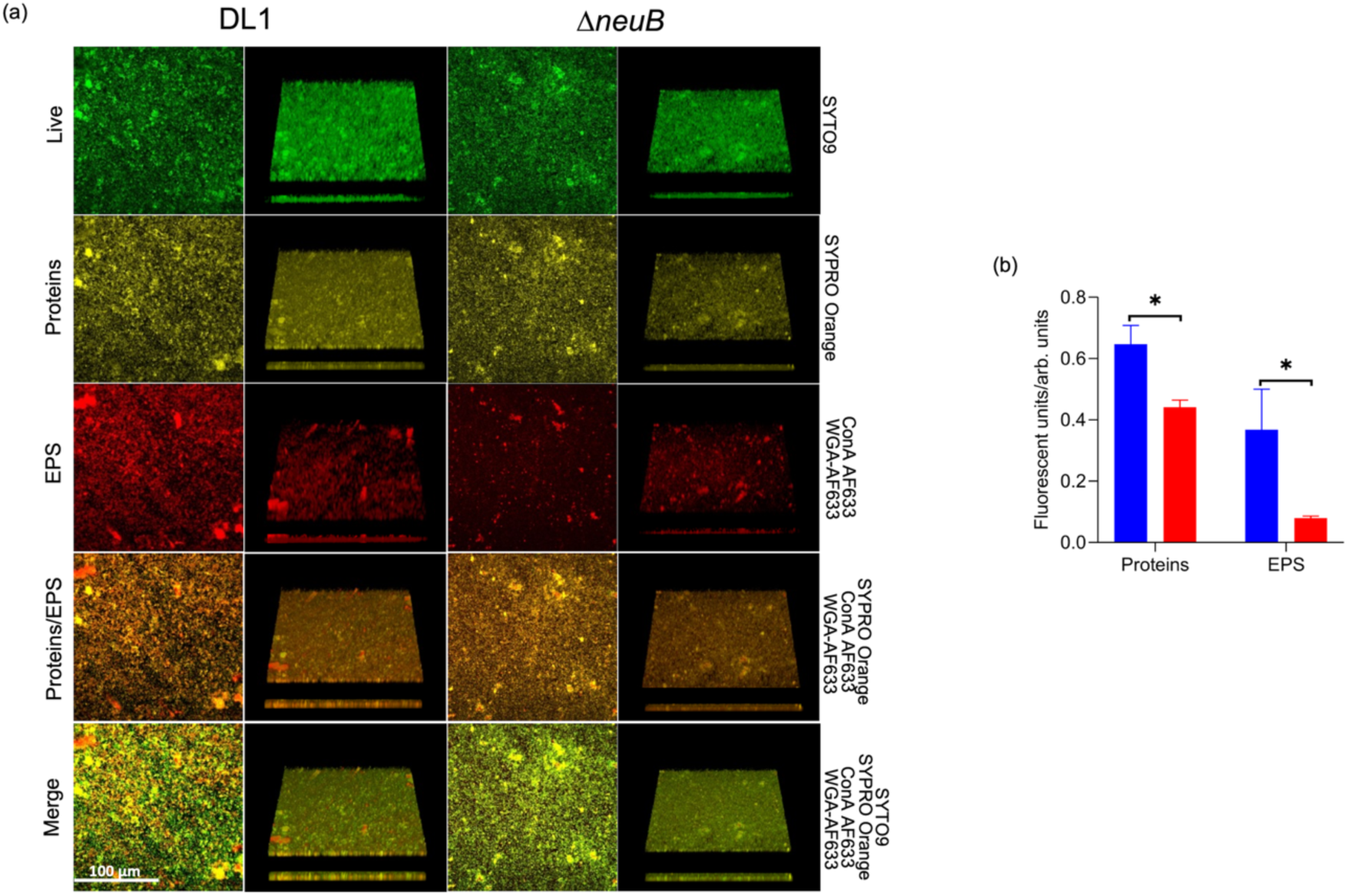
Detection of the principal components of *G. sulfurreducens* biofilms grown on hematite (Fe_2_O_3_-FTO) by CLSM. Analysis of biofilms acclimated by serial transfer for 2 weeks and grown on hematite-coated FTO (Fe_2_O_3_-FTO) electrodes in acetate-fumarate (NBAF) medium for 72 h. **(a)** Representative CLSM images of DL1 and Δ*neuB* biofilms stained with SYTO9 (green, viable cells), ConA/WGA-AF633 (red, extracellular polymeric substances [EPS]), and SYPRO Orange (orange, extracellular proteins). Scale bar = 100 µm. **(b)** Quantification of SYPRO Orange and ConA/WGA-AF633 fluorescence in DL1 and Δ*neuB* biofilms. Fluorescence intensity was expressed in arbitrary units for a biofilm volume of 4 × 10^5^ μm^3^ and normalized to SYTO9 fluorescence. Blue bars represent fluorescence signals from DL1 biofilms, whereas red bars represent fluorescence signals from Δ*neuB* biofilms. Bars indicate the mean of two independent experiments, and error bars represent standard deviation. Statistical significance was assessed using two-way ANOVA followed by Fisher’s least significant difference (LSD) test for pairwise comparisons without correction for multiple comparisons (*P* < 0.05; ns, not significant)

### Effect of the Δ*neuB* mutation on biofilm architecture on FTO electrodes

Biofilm development on bare FTO electrodes was also examined by CLSM (Fig. 6). Wild-type DL1 formed dense and homogeneous biofilms with an average thickness of 20.0 ± 3.1 μm. The Δ*neuB* mutant produced biofilms of similar thickness (18.0 ± 2.2 μm) (Fig. 6a). Although overall biofilm thickness remained largely unchanged, fluorescence imaging demonstrated a pronounced reduction in extracellular matrix abundance in the mutant. Both EPS-associated fluorescence and extracellular protein staining were significantly lower in Δ*neuB* biofilms than in the wild type (Fig. 6b,c), indicating substantial alterations in matrix composition.

**Fig. 6.**
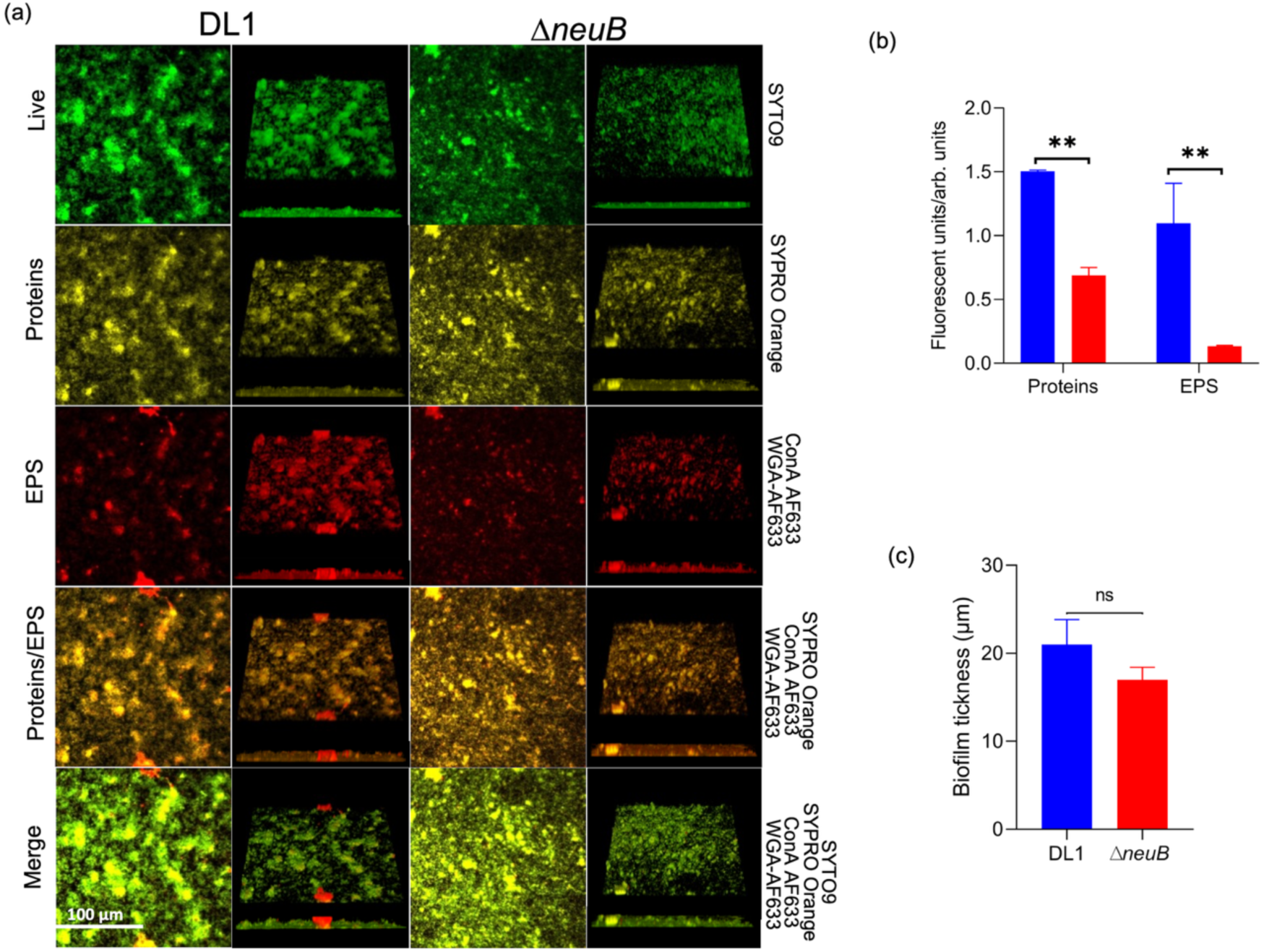
Detection of the principal components of *G. sulfurreducens* biofilms grown on FTO electrodes by CLSM. Analysis of biofilms acclimated by serial transfer for 2 weeks and grown on bare FTO electrodes in acetate-fumarate (NBAF) medium for 48 h. **(a)** Representative CLSM images of DL1 and Δ*neuB* biofilms stained with SYTO9 (green, viable cells), ConA/WGA-AF633 (red, extracellular polymeric substances [EPS]), and SYPRO Orange (orange, extracellular proteins). Scale bar = 100 µm. **(b)** Quantification of SYPRO Orange and ConA/WGA-AF633 fluorescence in DL1 and Δ*neuB* biofilms. Fluorescence intensity was expressed in arbitrary units for a biofilm volume of 4 × 10^5^ μm^3^ and normalized to SYTO9 fluorescence. Blue bars represent fluorescence signals from DL1 biofilms, whereas red bars represent fluorescence signals from Δ*neuB* biofilms. **(c)** Quantification of the thickness of biofilms formed by DL1 and Δ*neuB* strains. In **(b)** and **(c)**, bars indicate the mean of two independent experiments, and error bars represent standard deviation. Statistical significance was assessed using two-way ANOVA followed by Fisher’s least significant difference (LSD) test for pairwise comparisons without correction for multiple comparisons (ns, *P* > 0.05; * *P* < 0.05; ** *P* < 0.01)

### Electrochemical characterization of Δ*neuB* biofilms

The electrochemical properties of 48-h biofilms formed on FTO electrodes were evaluated using OCP, CV, and SWV (Fig. 7). OCP measurements showed that Δ*neuB* biofilms consistently exhibited more negative equilibrium potentials than wild-type biofilms, regardless of whether acetate was present (Fig. 7a). CV analysis further demonstrated enhanced electrochemical performance in the mutant. Under steady-state conditions, Δ*neuB* biofilms generated approximately 3-fold higher current densities than the wild type (Fig. 7b). Voltammetric data were fitting using a Nerst-Monod/Butler-Volmer-based kinetic model (R^2^>0.98). The non-linear regression indicated a shift in formal potential from approximately −0.107 V in the wild type to −0.134 V in the mutant, together with an increase in exchange current density from approximately 2.5 to 4.7 μA cm^−2^. SWV resolved distinct redox profiles for each strain (Fig. 7c,d). Wild-type biofilms displayed major peaks near −0.508, −0.164, and −0.027 V (vs. SHE), corresponding to the multi-cytochrome electron transport cascade spanning from periplasmic to outer-membrane cytochromes. In contrast, the Δ*neuB* mutant exhibited peaks centered at approximately −0.347, −0.272, and −0.085 V. Rather than assigning individual peaks to specific cytochromes, these potential shifts are interpreted as reflecting an altered local redox microenvironment within the EPS matrix, likely associated with differences in matrix density, electrostatic properties, and altered outer-membrane cytochrome organization.

**Fig. 7.**
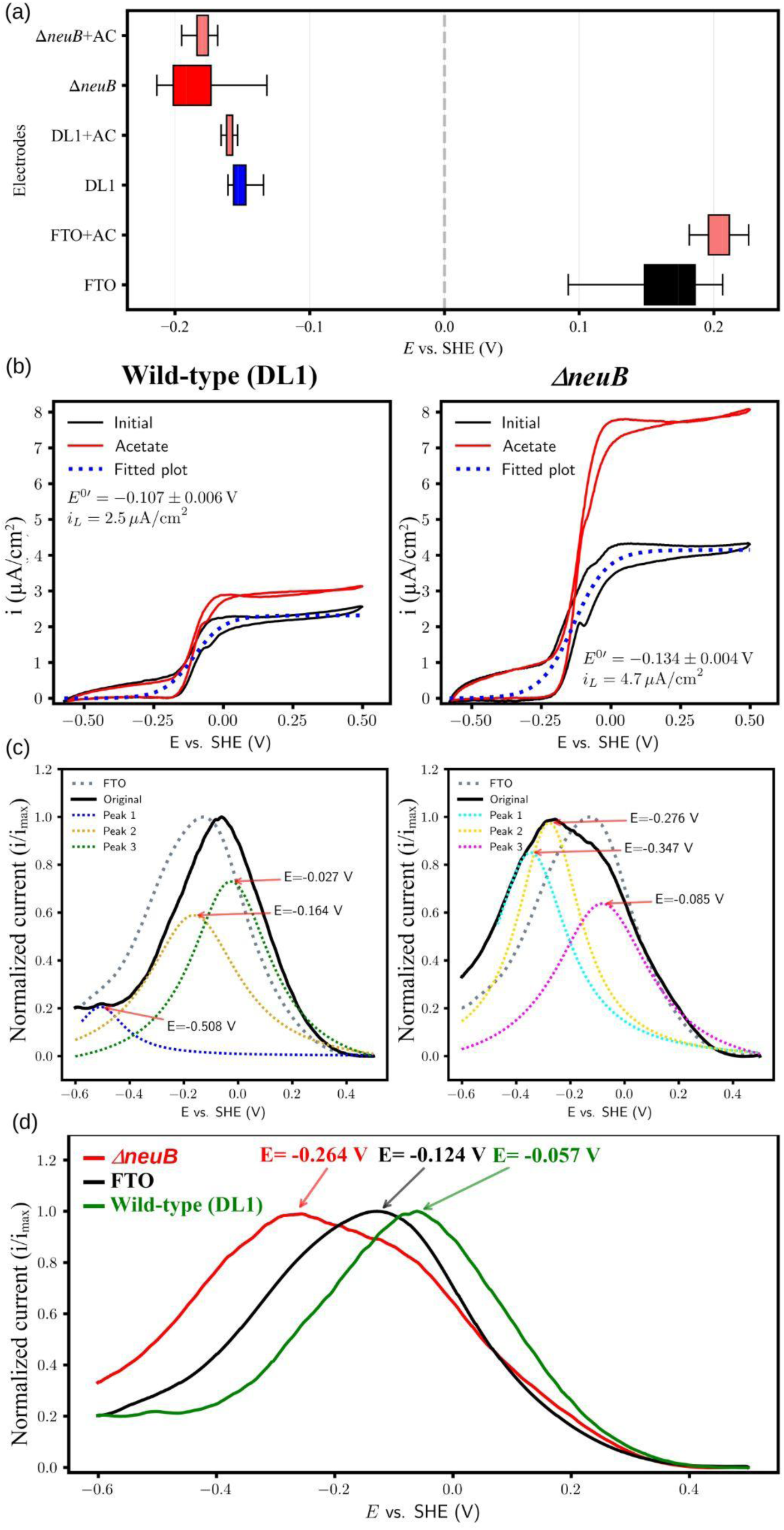
Electrochemical characterization of Δ*neuB* and DL1 biofilms grown on FTO electrodes. Characterization of 48-h biofilms acclimated by serial transfer for 2 weeks and grown on bare FTO electrodes in acetate-fumarate (NBAF) medium. **(a)** OCP measurements. **(b)** CV analysis; dashed lines represent the voltammetric fits, and the calculated formal potential (E*) values are indicated in each panel. **(c)** SWV analysis; representative profiles and peak deconvolution/fitting of the electrochemical responses of DL1 and Δ*neuB* strains are shown, with the E* values of individual peaks indicated. The responses were taken under the presence of sodium acetate (Turnover conditions) **(d)** Superimposition of SWV curves obtained for DL1 and Δ*neuB* biofilms and the bare FTO electrode without biofilm. Electrochemical measurements were performed in a three-electrode cell containing basal medium before (initial) and after the addition of acetate (20 mM) as an electron donor. All measurements were conducted under anaerobic conditions with continuous N_2_/CO_2_ sparging. In all panels, potentials are reported versus the standard hydrogen electrode (SHE)

The contrasting electrochemical behavior observed on insoluble Fe(III) oxides versus conductive FTO electrodes highlights the dual effect of matrix modification on extracellular electron transfer. While the altered, less dense EPS matrix in the Δ*neuB* mutant limits spatial access and electron transport toward insoluble, non-conductive iron minerals, it appears to reduce physical diffusion barriers on flat conductive surfaces. This reduced spatial constraint brings outer-membrane redox proteins into closer physical proximity to the FTO anode, facilitating direct interfacial electrical coupling. When combined with the selective upregulation of *omcS*, this enhanced interfacial contact accounts for the ∼3-fold increase in current density, the shift to a more negative formal potential, and the altered voltammetric profile. Similar phenomena have been described in *Shewanella oneidensis* MR-1, where EPS removal promoted direct contact between outer-membrane cytochromes and electrode surfaces, resulting in 2- to 3-fold higher current production (Gao et al. 2019).

### Expression of extracellular electron transfer genes in FTO-grown biofilms

To investigate transcriptional changes associated with enhanced electrochemical performance, expression of extracellular electron transfer genes was examined in biofilms grown on FTO electrodes (Fig. 8). Transcript levels of *omcB, omcC, omcE, omcS, omcZ, omcM, omcF, pgcA*, and *pilA* were quantified by RT-qPCR (Mehta et al. 2005; Reguera et al. 2005, 2007; Holmes et al. 2006; Inoue et al. 2010; Lovley et al. 2011; Qian et al. 2011; Liu et al. 2014; Zacharoff et al. 2017; Jiang et al. 2023). Expression of *omcZ* and *pilA* did not differ significantly between the wild-type and mutant strains. In contrast, *omcS* expression increased approximately 2.5-fold in the Δ*neuB* mutant relative to the wild type (Fig. 8), whereas no substantial differences were observed for the remaining genes analyzed.

**Fig. 8.**
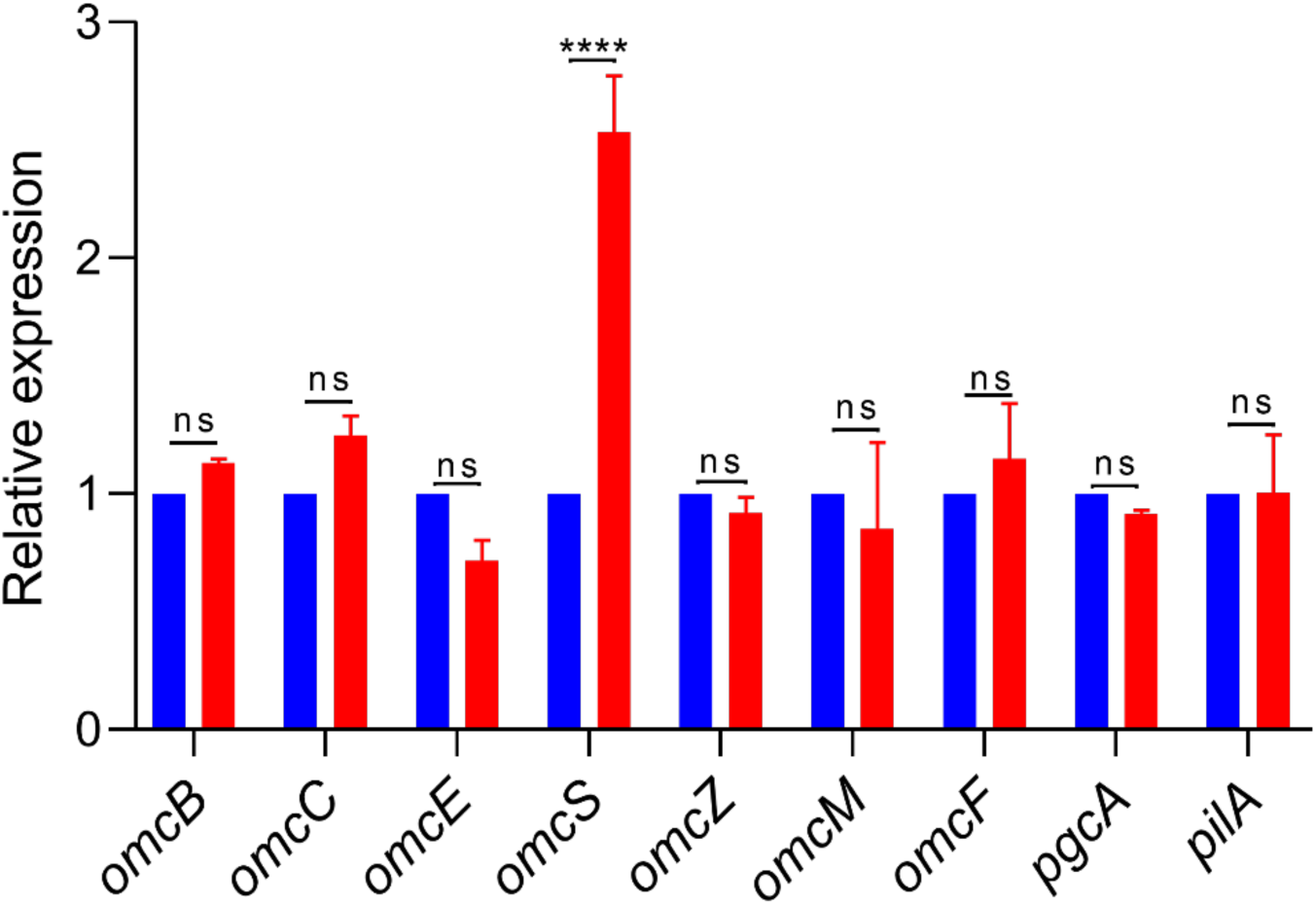
Analysis of gene expression in biofilms grown on FTO electrodes. Analysis of gene expression in *G. sulfurreducens* DL1 and Δ*neuB* biofilms acclimated by serial transfer for 2 weeks and grown on bare FTO electrodes in acetate-fumarate (NBAF) medium. The relative expression levels of genes encoding the cytochromes OmcB, OmcC, OmcE, OmcS, OmcZ, OmcM, OmcF, and PgcA, as well as the pilin subunit PilA, are shown. Blue bars represent gene expression levels in the wild-type DL1 strain, whereas red bars represent expression levels in the Δ*neuB* mutant strain. Error bars indicate the standard deviation of two independent experiments, and each experiment has technical triplicates per gene. Statistical significance was assessed using independent unpaired *t*-tests for each comparison without correction for multiple comparisons (ns, *P* > 0.05; * *P* < 0.05; ** *P* < 0.01; *** *P* < 0.001; **** *P* < 0.0001)

## Discussion

This study demonstrates that *neuB* (*gsu1970*) gene, putatively involved in nonulosonic acid biosynthesis, contributes to the extracellular matrix composition and functional properties of *G. sulfurreducens* biofilms. By deleting *neuB*, we established that disruption of this predicted biosynthetic pathway markedly alters WGA-reactive matrix glycoconjugates (Fig. 1), severely impairs insoluble Fe(III) oxide reduction (Fig. 3c), and associated with enhanced current production on FTO electrodes (Fig. 7). These findings indicate that *neuB*-dependent matrix components influence biofilm function in a substrate-dependent manner, highlighting distinct extracellular matrix requirements for electron transfer to mineral oxides versus conductive electrodes.

Sequence alignment and AlphaFold structural analyses support the designation of GSU1970 as a putative NeuB homolog in *G. sulfurreducens* (Supplementary Fig. 4). Although the predicted NeuB protein shares moderate sequence identity with homologs in *E. coli* and *N. meningitidis*, it retains 15 of the 16 conserved catalytic and metal-binding residues associated with NeuB activity (Supplementary Fig. 4). Our findings build upon earlier observations by Cologgi et al. (2021), who suggested that *gsu1970* participates in biofilm formation. By generating a scarless Δ*neuB* mutant and performing genetic complementation, we demonstrate that *gsu1970* is required for maintaining normal levels of WGA-reactive matrix glycoconjugates (Fig. 1a,b). The marked reduction in WGA-AF633 fluorescence observed in the Δ*neuB* mutant and its restoration upon complementation with pRneuB (Fig. 1a,c) confirms that these matrix alterations stem specifically from deletion of *gsu1970* rather than polar effects. Nevertheless, while this genetic evidence links *gsu1970* to matrix composition, we acknowledge that in silico modeling does not substitute for direct biochemical verification, such as in vitro enzymatic assays or LC–MS identification of Neu5Ac from extracted matrix fractions. Future analytical studies will be essential to definitively determine the chemical identity of the synthesized nonulosonic acid.

Loss of Neu5Ac also reduced biofilm viability, suggesting that this sugar contributes to the stability of the extracellular environment surrounding embedded cells. Similar protective functions have been reported in pathogenic bacteria, where surface-associated sialic acids promote biofilm persistence, reduce environmental stress, and enhance cell survival (Johnston et al. 2008). Although the ecological context differs considerably, our observations suggest that Neu5Ac performs a comparable structural role in *G. sulfurreducens*, helping to maintain a stable extracellular matrix that supports cell viability within electroactive biofilms. One of the most striking phenotypes of the Δ*neuB* mutant was its dramatic loss of insoluble Fe(III) oxide reduction while maintaining normal growth with soluble electron acceptors. Because respiration of insoluble metal oxides requires close contact between cells and the mineral surface, this phenotype suggests that Neu5Ac is specifically required for extracellular electron transfer to solid electron acceptors rather than for central metabolism or intracellular respiratory pathways. Previous studies have similarly demonstrated the importance of extracellular matrix organization for metal respiration in *G. sulfurreducens*. The *xap* locus (*gsu1498–gsu1508*) has been shown to participate in EPS matrix assembly, and deletion of *xapD* (*gsu1501*) markedly reduces cell aggregation, surface adhesion, and insoluble Fe(III) oxide reduction (Rollefson et al. 2011; Zhuang et al. 2020; Zhuang et al. 2022). These defects have been attributed to disruption of the extracellular matrix, which compromises the proper retention and organization of extracellular components required for efficient electron transfer. The similarities between the Δ*xapD* and Δ*neuB* phenotypes suggest that Neu5Ac contributes to the structural organization of the EPS matrix rather than directly participating in electron transfer reactions.

Several observations support this interpretation. UV-Vis analysis showed that extracellular *c*-type cytochromes remained present in the extracted EPS fraction (Supplementary Fig. 5), and RT-qPCR revealed no major downregulation of core EET genes during growth on hematite (Fig. 4). Together, these findings indicate that the severe defect in insoluble Fe(III) reduction is unlikely to stem from a transcriptional failure of the EET machinery. Instead, we hypothesize that disruption of the neuB-dependent glycan network compromises the matrix density (Fig. 5), which may indirectly impair the spatial orientation, physical packing, or surface accessibility of outer-membrane cytochromes relative to insoluble mineral acceptors. CLSM analysis demonstrated a marked reduction in EPS abundance in the Δ*neuB* mutant, indicating that *neuB*-dependent pathway contributes to overall extracellular matrix. We therefore propose that disruption of this predicted glycan network compromises matrix density and organization without altering overall biofilm thickness, thereby reducing the efficiency with which electroactive proteins interact with insoluble Fe(III) oxides.

This interpretation is consistent with previous studies of the *xap* locus, which have shown that disruption of EPS organization severely compromises Fe(III) oxide reduction despite relatively normal production of extracellular electron transfer proteins (Rollefson et al. 2011; Zhuang et al. 2020; Zhuang et al. 2022). Collectively, these observations suggest that efficient metal respiration depends not only on the presence of electroactive proteins but also on the structural organization of the extracellular matrix that positions these proteins at the cell-mineral interface. Although sialic acids have rarely been investigated in environmental biofilms, their occurrence in biofilms formed by the iron- and sulfur-oxidizing archaeon *Ferroplasma acidiphilum* further supports the possibility that nonulosonic acids contribute to biofilm organization in microorganisms that interact with mineral surfaces (Zhang et al. 2015). Our findings extend this concept by providing functional evidence that *neuB*-dependent matrix modification influences EPS matrix density and functional properties of electroactive biofilms involved in extracellular electron transfer.

In contrast to mineral reduction, the Δ*neuB* mutant exhibited enhanced electrochemical performance on FTO electrodes. Despite producing biofilms with reduced EPS and extracellular protein content, the mutant generated approximately 3-fold higher current densities, exhibited a more negative open-circuit potential, and showed substantial changes in its voltammetric profile (Fig. 7). These results indicate that a reduction in matrix abundance does not necessarily diminish electron transfer to conductive electrodes and may instead alter the physical organization of the biofilm in ways that facilitate electrode respiration. One possible explanation is that a less dense extracellular matrix decreases diffusion barriers or shortens the effective distance between electroactive proteins and the electrode surface. These structural changes could improve electrical coupling between cells and the anode, resulting in faster interfacial electron transfer. Consistent with this possibility, the Δ*neuB* mutant exhibited both a more negative formal potential and a higher exchange current density than the wild-type strain, indicating more efficient charge-transfer kinetics at the biofilm-electrode interface (LaBelle and Bond 2010; Snider et al. 2012; Elgrishi et al. 2018; Istanbullu et al. 2019; Schrott et al. 2019).

The distinct SWV profiles observed for the two strains further support the conclusion that deletion of *neuB* alters the electrochemical organization of the biofilm (Fig. 7d) (Istanbullu et al. 2019). Shifts in peak position likely reflect changes in the accessibility, relative contribution, or microenvironment of extracellular redox components rather than the appearance of entirely new electron transfer pathways. Although the identities of the individual redox centers cannot be assigned solely from voltammetric data, the reproducible changes in the electrochemical fingerprints demonstrate that Neu5Ac deficiency substantially remodels the redox architecture of the biofilm (Mirčeski et al. 2007; Istanbullu et al. 2019). The transcriptional response observed during growth on FTO electrodes provides additional insight into this phenotype. Unlike biofilms grown on hematite, the Δ*neuB* mutant exhibited a pronounced increase in *omcS* expression, whereas the expression of *omcZ* and *pilA* remained largely unchanged (Fig. 8).

OmcS has been shown to facilitate extracellular electron transfer, particularly during the early stages of biofilm development and in relatively thin biofilms, where alternative long-range conductive network architectures may not yet be fully established (Nevin et al. 2009; Holmes et al. 2006; Wang et al. 2022). Structural studies further indicate that OmcS can decorate conductive pili or assemble into extracellular filaments that promote direct electrical coupling with solid surfaces (Leang et al. 2010; Steidl et al. 2016; Lovley and Walker 2019; Filman et al. 2019; Neu et al. 2022; Portela et al. 2024; Wang et al. 2022). In our study, CLSM analysis showed that Δ*neuB* biofilms developed a significantly less dense EPS matrix on FTO electrodes. This physical alteration may simulate a spatially unconstrained, early-stage biofilm environment where cells rely heavily on OmcS-mediated interfacial electron transfer to the anode. Consequently, the selective upregulation of *omcS* (∼2.5-fold) observed in FTO-grown Δ*neuB* biofilms likely represents an adaptive compensatory response to overcome structural matrix deficits. A similar phenomenon has been reported in the Δ*gsu1771* mutant, where *omcS* overexpression accompanied enhanced current generation (Hernández-Eligio et al. 2022; Rodríguez-Torres et al. 2025). Although the precise mechanism remains to be determined, our results suggest that disruption of *neuB*-mediated glycan biosynthesis triggers a compensatory remodeling of the extracellular electron transfer network through selective upregulation of *omcS*. This adaptive response may contribute to the more negative electrochemical potential and enhanced current production observed in the Δ*neuB* mutant by improving electron transfer across thin, electrode-associated biofilms. Interestingly, this compensatory response was observed only during growth on FTO electrodes and not on hematite, indicating that matrix remodeling depends on the physical and chemical nature of the terminal electron acceptor.

Interestingly, this compensatory response was observed only during growth on FTO electrodes and not during growth on hematite, indicating that matrix remodeling depends on the nature of the extracellular electron acceptor. These findings suggest that *G. sulfurreducens* employs distinct adaptive strategies for electron transfer to conductive electrodes and insoluble metal oxides, even though both processes rely on overlapping extracellular electron transfer machinery. Overall, our results support a model in which the *neuB* gene contributes to EPS matrix composition and density rather than as a direct regulator of extracellular electron transfer gene expression. During Fe(III) oxide reduction, Neu5Ac promotes the organization of an extracellular matrix that enables effective electrical coupling between cells and mineral surfaces. In contrast, its absence results in a less complex matrix that appears to favor electron transfer to conductive electrodes while simultaneously triggering increased *omcS* expression. These findings establish *neuB*-dependent glycoconjugates as important determinants of biofilm architecture and demonstrate that matrix composition can differentially influence extracellular electron transfer depending on the nature of the terminal electron acceptor.

## Conclusions

In conclusion, this study demonstrates that the *neuB* gene (*gsu1970*) is required for maintaining normal levels of WGA-reactive glycoconjugates in the extracellular matrix of *Geobacter sulfurreducens*. Deletion of *neuB* did not affect growth with soluble fumarate or ferric citrate but reduced insoluble Fe(III) oxide reduction by approximately 91% (Fig. 3). This reduction was not accompanied by significant downregulation of core EET genes (Fig. 4), suggesting that the impaired Fe(III) reduction is associated with alterations in EPS matrix density rather than transcription suppression of electron transfer proteins. Conversely, the Δ*neuB* mutant exhibited enhanced current density on FTO electrodes, a shift in formal midpoint potential, and elevated *omcS* transcriptional levels (Fig. 7, Fig. 8). Collectively, these findings show that *neuB*-dependent matrix modification alters EPS matrix density without disrupting gross 3D biofilm thickness, differentially affects electron transfer depending on the terminal electron acceptor and highlight the role of carbohydrate biosynthetic pathways in modulating electroactive biofilm.

## Acknowledgments

The authors express their gratitude to Andrés Saralegui, Maricela Olvera, Shirley Ainsworth, and Leticia Olvera for their excellent technical assistance. BJ-R wishes to thank the Secretaría de Ciencia, Humanidades, Tecnología e Innovación (SECIHTI) for the doctoral fellowships provided. We are also grateful to Dalia Alejandra Mazón-Montijo for kindly providing the Fe_2_O_3_-FTO electrodes, and to Gloria Soberón and Ricardo Oropeza for their insightful comments and valuable suggestions which greatly improve the quality of this manuscript.

## Contributions

Bernardo Jaramillo-Rodríguez: conceptualization and design, data analysis and visualization, writing-original draft preparation, writing-review and editing. Eder Gómez-Escorza: data analysis and visualization. Guillermo Huerta-Miranda: data analysis and visualization, writing-original draft preparation, writing-review and editing. Katy Juárez: conceptualization and design, data analysis and visualization, writing-original draft preparation, writing-review and editing. Alberto Hernández-Eligio: conceptualization and design, data analysis and visualization, writing-original draft preparation, writing-review and editing. All authors read, critically revised, and approved the final version of the manuscript.

## Funding

AH-E, received financial support through the Secretaría de Ciencia, Humanidades, Tecnología e Innovación (SECIHTI, Grant No. CBF2023-2024-726). KJ received funding from the Programa de Apoyos a Proyectos de Investigación e Innovación Tecnológica (PAPIIT, Universidad Nacional Autónoma de México, Grant No. IN212325).

## Ethics declarations

### Ethics approval

This article does not contain any studies involving human participants or animals performed by any of the authors.

### Competing interests

The authors declare no competing interests.

## Data availability statements

The authors declare that the data supporting the findings of this study are available within the paper and its Supplementary Information file. All primer sequences, bacterial strains, and plasmids are provided in Supplementary Table 1, along with original reference used in this study.

## Notes

### Competing Interest Statement

The authors have declared no competing interest.

## References

Almagro-Moreno S, Boyd EF (2009) Insights into the evolution of sialic acid catabolism among bacteria. BMC Evol Biol 9:118. doi:10.1186/1471-2148-9-118

Boleij M, Kleikamp H, Pabst M, Neu TR, van Loosdrecht MCM, Lin Y (2020) Decorating the Anammox House: Sialic acids and sulfated glycosaminoglycans in the extracellular polymeric substances of Anammox granular sludge. Environ Sci Technol 54(8):5218–5226. doi:10.1021/acs.est.9b07207

Bond DR, Lovley DR (2003) Electricity production by *Geobacter sulfurreducens* attached to electrodes. Appl Environ Microbiol 69(3):1548–1555. doi:10.1128/AEM.69.3.1548-1555.2003

Cao B, Shi L, Brown RN, Xiong Y, Fredrickson JK, Romine MF, Marshall MJ, Lipton MS, Beyenal H (2011) Extracellular polymeric substances from *Shewanella* sp. HRCR-1 biofilms: characterization by infrared spectroscopy and proteomics. Environ Microbiol 13(4):1018–1031. doi:10.1111/j.1462-2920.2010.02407.x

Cardona L, Saini JS, Rodilla Ramírez PN, Adler A, Holliger C (2025) Multiple extracellular polymeric substance pathways transcribed by *Accumulibacter* and the flanking community during aerobic granule formation and after influent modification. Appl Environ Microbiol 91(4):e0176924. doi:10.1128/aem.01769-24

Chan CH, Levar CE, Zacharoff L, Badalamenti JP, Bond DR (2015) Scarless genome editing and stable inducible expression vectors for Geobacter sulfurreducens. Appl Environ Microbiol 81(20):7178–7186. doi:10.1128/AEM.01967-15

Chen LM, de Bruin S, Pronk M, Sousa DZ, van Loosdrecht MCM, Lin Y (2023) Sialylation and sulfation of anionic glycoconjugates are common in the extracellular polymeric substances of both aerobic and anaerobic granular sludges. Environ Sci Technol 57(35):13217–13225. doi:10.1021/acs.est.2c09586

Cholula-Calixto J, Huerta-Miranda G, Jaramillo-Rodríguez B, Bustamante VH, Juarez K, Hernández-Eligio A (2026) FliW regulates biofilm formation in *Geobacter sulfurreducens* through interaction with CsrA. App Microbiol Biotech, 110:223. doi: 10.1007/s00253-026-13884-0

Cologgi DL, Otwell AE, Speers AM, Rotondo JA, Reguera G (2021) Genetic analysis of electroactive biofilms. Int Microbiol 24:631–648. doi:10.1007/s10123-021-00176-y

Coppi MV, Leang C, Sandler SJ, Lovley DR (2001) Development of a genetic system for *Geobacter sulfurreducens*. Appl Environ Microbiol 67:3180–3187. doi:10.1128/AEM.67.7.3180-3187.2001

Cummings EA, Mailley P, Linquette-Mailley S, Eggins BR, McAdams ET, McFadden S (1998) Amperometric carbon paste biosensor based on plant tissue for the determination of total flavanol content in beers. Analyst, 123(10):1975–1980. doi:10.1039/a804021d

Dhar BR, Ryu H, Ren H, Domingo JWS, Chae J, Lee H (2016) High Biofilm Conductivity Maintained Despite Anode Potential Changes in a Geobacter-Enriched Biofilm. ChemSusChem, 9(24):3485–3491. doi:10.1002/cssc.201601007

Elgrishi N, Rountree KJ, McCarthy BD, Rountree ES, Eisenhart TT, Dempsey JL (2018) A practical beginner’s guide to cyclic voltammetry. J Chem Educ 95:197–206. doi:10.1021/acs.jchemed.7b00361

Filman DJ, Marino SF, Ward JE, Yang L, Mester Z, Bullitt E, Lovley DR, Strauss M (2019) Cryo-EM reveals the structural basis of long-range electron transport in a cytochrome-based bacterial nanowire. Commun Biol 2:219. doi:10.1038/s42003-019-0448-9

Gao L, Lu X, Liu H, Li J, Li W, Song R, Wang R, Zhang D, Zhu J (2019) Mediation of Extracellular Polymeric Substances in Microbial Reduction of Hematite by *Shewanella oneidensis* MR-1. Front Microbiol 10:1–12. doi:10.3389/fmicb.2019.00575

Ghosh S (2020) Sialic acid and biology of life: An introduction. Sialic Acids and Sialoglyconjugates in the Biology of Life, Health and Disease In: Elsevier eBooks, pp 1–61. doi:10.1016/b978-0-12-816126-5.00001-9

Gunawan J, Simard D, Gilbert M, Lovering AL, Wakarchuk WW, Tanner ME, Strynadka NCJ (2005) Structural and Mechanistic Analysis of Sialic Acid Synthase NeuB from *Neisseria meningitidis* in Complex with Mn2+, Phosphoenolpyruvate, and N-Acetylmannosaminitol. J Biol Chem 280(5):3555–3563. doi:10.1074/jbc.M411942200

Hernández-Eligio A, Huerta-Miranda GA, Martínez-Bahena S, Castrejón-López B, Miranda-Hernández M, Juárez K (2022) GSU1771 regulates extracellular electron transfer and electroactive biofilm formation in *Geobacter sulfurreducens*: Genetic and electrochemical characterization. Bioelectrochem 145:108101. doi:10.1016/j.bioelechem.2022.108101

Holmes DE, Chaudhuri SK, Nevin KP, Mehta T, Methé BA, Liu A, Ward JE, Woodard TL, Webster J, Lovley DR (2006) Microarray and genetic analysis of electron transfer to electrodes in *Geobacter sulfurreducens*. Environ Microbiol 8(10):1805–1815. 10.1111/j.1462-2920.2006.01065.x

Huerta-Miranda GA, Arroyo-Escoto AI, Burgos X, Juárez K, Miranda-Hernández M (2019) Influence of the major pilA transcriptional regulator in electrochemical responses of *Geobacter sulfurreducens* PilR-deficient mutant biofilm formed on FTO electrodes. Bioelectrochem 127:145–153. doi:10.1016/j.bioelechem.2019.02.006

Huerta-Miranda GA, Rodríguez-Torres LM, Martínez-García AL, Mazón-Montijo DA, Hernández-Eligio A, Miranda-Hernández M, Juárez K (2023) *Geobacter sulfurreducens* electroactive biofilms on Fe_2_O_3_/FTO support-electrodes for developing a sodium acetate electrochemical biosensor. Biosens Bioelectron: X, 14:100370. doi:10.1016/j.biosx.2023.100370

Inoue K, Qian X, Morgado L, Kim B-C, Mester T, Izallalen M, Salgueiro CA, Lovley DR (2010) Purification and characterization of OmcZ, an outer-surface, multiheme c-type cytochrome essential for optimal current production by *Geobacter sulfurreducens*. Appl Environ Microbiol 76(12):3999–4007. 10.1128/AEM.00027-10

Istanbullu O, Babauta J, Renslow R, Beyenal H (2019) Monitoring Electron Transfer Rates of Electrode-Respiring Cells. In: Tiquia-Arashiro SM, Pant D (eds) Microbial electrochemical technologies. CRC Press. pp.76–84. doi:10.1201/9780429487118

Jaramillo-Rodríguez JB, Vega-Alvarado L, Rodríguez-Torres LM, Huerta-Miranda GA, Hernández-Eligio A, Juarez K (2023) Global transcriptional analysis of *Geobacter sulfurreducens gsu1771* mutant biofilm grown on two different support structures. PLOS ONE 18(10):e0293359. doi:10.1371/journal.pone.0293359

Jiang J, He P, Luo Y, Peng Z, Jiang Y, Hu Y, Qi L, Dong X, Dong Y, Shi L (2023) The varied roles of *pilA*-N, *omcE*, *omcS*, *omcT*, and *omcZ* in extracellular electron transfer by *Geobacter sulfurreducens*. Front Microbiol 14:1251346. doi:10.3389/fmicb.2023.1251346

Johnston WJ, Coussens NP, Allen S, Houtman JCD, Turner KH, Zaleski A, Ramaswamy S, Gibson BW, Apicella MA (2008) Characterization of the N-acetyl-5-neuraminic acid-binding site. J Biol Chem 283(2):855–865. doi:10.1074/jbc.M706603200

Jones GD, Jones MG, Wilson MT, Brunori M, Colosimo A, Sarti P (1983) Reactions of cytochrome c oxidase with sodium dithionite. Biochem J 209(1):175–182. doi:10.1042/bj2090175

Kim BC, Leang C, Ding YH, Glaven RH, Coppi MV, Lovley DR (2005) OmcF, a putative c-type monoheme outer membrane cytochrome required for the expression of other outer membrane cytochromes in *Geobacter sulfurreducens*. J Bacteriol 187(13):4505–4513. doi:10.1128/JB.187.13.4505-4513.2005

Kleikamp HBC, Lin YM, McMillan DGG, Geelhoed JS, Naus-Wiezer SNH, van Baarlen P, Saha C, Louwen R, Sorokin DY, van Loosdrecht MCM, Pabst M (2020) Tackling the chemical diversity of microbial nonulosonic acids - a universal large-scale survey approach. Chem Sci 11(11):3074–3080. doi:10.1039/c9sc06406k

LaBelle E, Bond DR (2010) Cyclic voltammetry for the study of microbial electron transfer at electrodes. In: Rabaey K, Angenent L, Schröder U, Keller J (ed) Bioelectrochemical Systems: From Extracellular Electron Transfer to Biotechnological Application. IWA, Publishing 137–152. doi: 10.2166/9781780401621

Leang C, Qian X, Mester T, Lovley DR (2010) Alignment of the c-type cytochrome OmcS along pili of *Geobacter sulfurreducens*. Appl Environ Microbiol 76(12):4080–4084. doi:10.1128/AEM.00023-10

Liu Y, Wang Z, Liu J, Levar C, Edwards MJ, Babauta JT, Kennedy DW, Shi Z, Beyenal H, Bond DR, Clarke TA, Butt JN, Richardson DJ, Rosso KM, Zachara JM, Fredrickson JK, Shi L (2014) A trans-outer membrane porin-cytochrome protein complex. Environ Microbiol Rep 6(6):776–785. 10.1111/1758-2229.12204

Lovley DR, Phillips EJP (1988) Novel mode of microbial energy metabolism: Organic carbon oxidation coupled to the reduction of iron or manganese. Appl Environ Microbiol 54(6):1472–1480. doi:10.1128/aem.54.6.1472-1480.1988

Lovley DR, Ueki T, Zhang T, Malvankar NS, Shrestha PM, Flanagan KA, Aklujkar M, Butler JE, Giloteaux L, Rotaru A-E, Holmes DE, Franks AE, Orellana R, Risso C, Nevin KP (2011) *Geobacter*: the microbe electric’s physiology, ecology, and practical applications. Adv Microb Physiol 59:1–100. doi:10.1016/B978-0-12-387661-4.00004-5

Lovley DR, Walker DJF (2019) *Geobacter* protein nanowires. Front Microbiol 10:2078. doi:10.3389/fmicb.2019.02078

Methé BA, Nelson KE, Eisen JA, Paulsen IT, Nelson W, Heidelberg JF, Wu D, Wu M, Ward N, Beanan MJ, Dodson RJ, Madupu R, Brinkac LM, Daugherty SC, DeBoy RT, Durkin AS, Gwinn M, Kolonay JF, Sullivan SA, Haft DH, Selengut J, Davidsen TM, Zafar N, White O, Tran B, Romero C, Forberger HA, Weidman J, Khouri H, Feldblyum TV, Utterback TR, Van Aken SE, Lovley DR, Fraser CM (2003) Genome of *Geobacter sulfurreducens*: metal reduction in subsurface environments. Science 302(5652):1967–1969. doi:10.1126/science.1088727

Marcus AK, Torres CI, Rittmann BE (2007) Conduction-based modeling of the biofilm anode of a microbial fuel cell. Biotechnol Bioeng, 98:1171–1182. doi:10.1002/bit.21533

Mehta T, Coppi MV, Childers SE, Lovley DR (2005) Outer membrane c-type cytochromes required for Fe(III) and Mn(IV) oxide reduction in *Geobacter sulfurreducens*. Appl Environ Microbiol 71(12):8634–8641. doi:10.1128/AEM.71.12.8634-8641.2005

Mirceski V, Komorsky-Lovric S, Lovric M (2007) Square-wave voltammetry: Theory and application. Springer Berlin, Heidelberg. doi:10.1007/978-3-540-73740-7

Nakamura R, Ishii K, Hashimoto K (2009) Electronic absorption spectra and redox properties of C type cytochromes in living microbes. Angew Chem Int Ed Engl 48:1606–1608. doi:10.1002/anie.200804917

Neu J, Shipps CC, Guberman-Pfeffer MJ, Shen C, Srikanth V, Spies JA, Kirchhofer ND, Yalcin SE, Brudvig GW, Batista VS, Malvankar NS (2022) Microbial biofilms as living photoconductors due to ultrafast electron transfer in cytochrome OmcS nanowires. Nat Commun 13:5150. doi:10.1038/s41467-022-32659-5

Nevin KP, Kim B-C, Glaven RH, Johnson JP, Woodard TL, Methé BA, Didonato Jr RJ, Covalla SF, Franks AE, Liu A, Lovley DR (2009) Anode biofilm transcriptomics reveals outer surface components required for maximal current production in *Geobacter sulfurreducens*. PLoS ONE 4(5):e5628. 10.1371/journal.pone.0005628

Paleček E, Tkáč J, Bartošík M, Bertók T, Ostatná V, Paleček J (2015) Electrochemistry of nonconjugated proteins and glycoproteins: Toward sensors for biomedicine and glycomics. Chem Rev 115(5):2045–2108. doi:10.1021/cr500279h

Petersen M, Fessner W-D, Frosch M, Lüneberg E (2000) The *siaA* gene involved in capsule polysaccharide biosynthesis of *Neisseria meningitidis* B codes for N-acylglucosamine-6-phosphate 2-epimerase activity. FEMS Microbiol Lett 184(2):161–164. doi:10.1111/j.1574-6968.2000.tb09008.x

Pinel ISM, Kleikamp HBC, Pabst M, Vrouwenvelder JS, van Loosdrecht MCM, Lin Y (2020) Sialic acids: An important family of carbohydrates overlooked in environmental biofilms. Appl Sci 10(21):7694. doi:10.3390/app10217694

Portela PC, Shipps CC, Shen C, Srikanth V, Salgueiro CA, Malvankar NS (2024) Widespread extracellular electron transfer pathways for charging microbial cytochrome OmcS nanowires via periplasmic cytochromes PpcABCDE. Nat Commun 15(1):2434. doi:10.1038/s41467-024-46192-0

Qian X, Mester T, Morgado L, Arakawa T, Sharma ML, Inoue K, Joseph C, Salgueiro CA, Maroney MJ, Lovely DR (2011) Biochemical characterization of purified OmcS, a c-type cytochrome required for insoluble Fe(III) reduction in *Geobacter sulfurreducens*. Biochim Biophys Acta 1807(4):404–412. doi:10.1016/j.bbabio.2011.01.003

Reguera G, McCarthy KD, Mehta T, Nicoll JS, Tuominen MT, Lovley DR (2005) Extracellular electron transfer via microbial nanowires. Nature 435(7045):1098–1101. doi:10.1038/nature03661

Reguera G, Pollina RB, Nicoll JS, Lovley DR (2007) Possible nonconductive role of *Geobacter sulfurreducens* pilus nanowires in biofilm formation. J Bacteriol 189(5):2125–2127. doi:10.1128/JB.01284-06

Reguera G, Kashefi K (2019) The electrifying physiology of *Geobacter* bacteria, 30 years on. Adv Microb Physiol 74:1–96. doi:10.1016/bs.ampbs.2019.02.007

Rodríguez-Torres LM, Huerta-Miranda GA, Martínez-García AL, Mazón-Montijo DA, Hernández-Eligio A, Miranda-Hernández M, Juárez K (2025) Influence of support materials on the electroactive behavior, structure and gene expression of wild type and GSU1771-deficient mutant of *Geobacter sulfurreducens biofilms*. Environ Sci Pollut Res Int 32(28):16740–16759. doi:10.1007/s11356-024-33612-3

Rollefson JB, Stephen CS, Tien M, Bond DR (2011) Identification of an extracellular polysaccharide network essential for cytochrome anchoring and biofilm formation in *Geobacter sulfurreducens*. J Bacteriol 193(5):1023–1033. doi:10.1128/JB.01092-10

Sambrook J, Fritsch EF, Maniatis T (1989) Molecular Cloning: A Laboratory Manual, 2nd edn. Cold Spring Harbor Laboratory Press, Cold Spring Harbor

Schindelin J, Arganda-Carreras I, Frise E, Kaynig V, Longair M, Pietzsch T, Preibisch S, Rueden C, Saalfeld S, Schmid B, Tinevez J-Y, White DJ, Hartenstein V, Eliceiri K, Tomancak P, Cardona A (2012) Fiji: An open-source platform for biological-image analysis. Nat Methods 9(7):676–682. doi:10.1038/nmeth.2019

Schrott G, Bonanni PS, Busalmen JP (2019) Open circuit potentiometry reports on internal redox states of cells in *Geobacter sulfurreducens* biofilms. Electrochim Acta 303:176–182. doi:10.1016/j.electacta.2019.02.078

Snider RM, Strycharz-Glaven SM, Tsoi SD, Erickson JS, Tender LM (2012) Long-range electron transport in *Geobacter sulfurreducens* biofilms is redox gradient-driven. Proc Natl Acad Sci USA 109(38):15467–15472. doi:10.1073/pnas.1209829109

Steidl RJ, Lampa-Pastirk S, Reguera G (2016) Mechanistic stratification in electroactive biofilms of *Geobacter sulfurreducens*. Nat Commun 7:12217. doi:10.1038/ncomms12217

Torres CI, Marcus AK, Parameswaran P, Rittmann BE (2008) Kinetic Experiments for Evaluating the Nernst−Monod Model for Anode-Respiring Bacteria (ARB) in a Biofilm Anode. Environ Sci Technol, 42(17):6593–6597. doi:10.1021/es800970w

Torres CI, Marcus AK, Lee H-S, Parameswaran P, Krajmalnik-Brown R, Rittmann BE (2010) A kinetic perspective on extracellular electron transfer by anode-respiring bacteria. FEMS Microbiol Rev, 34(1):3–17. doi:10.1111/j.1574-6976.2009.00191.x

Vann WF, Daines DA, Murkin AS, Tanner ME, Chaffin DO, Rubens CE, Vionnet J, Silver RP (2004) The NeuC protein of *Escherichia coli* K1 is a UDP N-acetylglucosamine 2-epimerase. J Bacteriol 186(3):706–712. doi:10.1128/JB.186.3.706-712.2004

Vieira AZ, Raittz RT, Faoro H (2021) Origin and evolution of nonulosonic acid synthases and their relationship with bacterial pathogenicity revealed by a large-scale phylogenetic analysis. Microb Genom 7(4):000563. doi:10.1099/mgen.0.000563

Wang Z, Hu Y, Dong Y, Shi L, Jiang Y (2023) Enhancing electrical outputs of the fuel cells with *Geobacter sulfurreducens* by overexpressing nanowire proteins. Microbial Biotechnology, 16:534–545. doi:10.1111/1751-7915.14128

Xu J, Zhou W, Han X, Liu J, Dong Y, Jiang Y, Zhong Y, Shi L, Hu Y (2025) Extracellular electron transfer proteins contribute to reduction of ferric minerals by *Geobacter* biofilms. Appl Environ Microbiol 91(5):e0036925. doi:10.1128/aem.00369-25

Zacharoff LA, Morrone DJ, Bond DR (2017) *Geobacter sulfurreducens* extracellular multiheme cytochrome PgcA facilitates respiration to Fe(III) oxides but not electrodes. Front Microbiol 8:2481. doi:10.3389/fmicb.2017.02481

Zhang RY, Neu TR, Bellenberg S, Kuhlicke U, Sand W, Vera M (2015) Use of lectins to in situ visualize glycoconjugates of extracellular polymeric substances in acidophilic archaeal biofilms. Microb Biotechnol 8(3):448–461. doi:10.1111/1751-7915.12188

Zhuang Z, Yang G, Mai Q, Guo J, Liu X, Zhuang L (2020) Physiological potential of extracellular polysaccharide in promoting *Geobacter* biofilm formation and extracellular electron transfer. Sci Total Environ 741:140365. doi:10.1016/j.scitotenv.2020.140365

Zhuang Z, Yang G, Zhuang L (2022) Exopolysaccharides matrix affects the process of extracellular electron transfer in electroactive biofilms. Sci Total Environ 806(3):150713. doi:10.1016/j.scitotenv.2021.150713

