## Supplementary Information for "Effect of *neuB* deletion on biofilm formation and extracellular electron transfer in *Geobacter sulfurreducens*"

BJ-R 0000-0002-4081-7440

Keywords: *Geobacter sulfurreducens*, biofilm, electrochemical analysis, N-acetylneuraminic acid synthesis, *neuB* gene

**Supplementary Table 1** List of strains, plasmids, and oligonucleotides used in this work.

| **Name** | **Description** | **Reference** |
| --- | --- | --- |
| **Strains** |  |  |
| *Geobacter sulfurreducens* |  |  |
| DL1 | *G. sulfurreducens* wild type strain | (Caccavo et al. 1994) |
| Δ*neuB* | DL1 strain with the *neuB* gene eliminated | This work |
| *Escherichia coli* |  |  |
| S17-1 | Strain used to perform the conjugation process | (Simon et al. 1983) |
| MC1061 | Strain used for plasmid maintenance and cloning transformation | (Casadaban y Cohen, 1980) |
| **Plasmids** |  |  |
| pJET1.2 | Linearized plasmid from the clonejet pcr cloning kit. | Thermo Scientific |
| pRRrecA | pJET1.2 plasmid with the *G. sulfurreducens recA* regulatory region (RRrecA) | This work |
| pRG5.1 | Plasmid used for complementation | (Kim et al. 2005) |
| pRRrecA-neuB | pJET1.2 plasmid with the RR*recA-neuB* fragment | This work |
| pRneuB | pRG5.1 plasmid with RR*recA-neuB* fragment | This work |
| pK18mobSacB | Suicide vector used to perform mutagenesis | (Chan et al. 2015) |
| pK18mobsacB-ΔneuB | Plasmid used for construct the Δ*neuB* mutant strain | This work |
| **Oligonucleotides** |  |  |
| *For mutant construction* |  |  |
| Primer1-HindIII-neuB | ATAAGCTTCGGATGATGTTCCATTCCAG | This work |
| Primer2-neuB | GGAAATGTTCCGGTTGGAGCGGACATGCAGGAAGATGACG | This work |
| Primer3-neuB | CGTCATCTTCCTGCATGTCCGCTCCAACCGGAACATTTCC | This work |
| Primer4-EcoRI-neuB | ATGAATTCGACCCATTAAGACTCTGGCG | This work |
| *For mutant complementation* |  |  |
| EcoRI-recAPromFw | ATGAATTCCCTGGCGGGAGAC | This work |
| NdeI-recAPromRv | CATATGCGCCTCCTGGGCGG | This work |
| NdeI-neuBFw | CATATGAACAGCATGAAATTAGG | This work |
| HindIII-neuBRv | AAGCTTTCACACCAGATCCTCCCAGC | This work |
| *for RTqPCR* |  |  |
| recA660f | GTGAAGGTGGTCAAGAACAAGGT | Holmes et al. 2006 |
| recA737r | GGAAATGCCCTCACCGTAGTAA | Holmes et al. 2006 |
| qPCRpilAfw | AATTACCCCCATACCCCAAC | Hernández-Eligio et al. 2020 |
| qPCRpilArev | AGCAGCTCGATAAGGGTGAA | Hernández-Eligio et al. 2020 |
| qPCRomcBfw | GGAGTATGTGGCATCCCTTG | Hernandez-Eligio et al. 2020 |
| qPCRomcBrev | ACCGTTGGCATTCGTATCTC | Hernandez-Eligio et al. 2020 |
| qPCRomcCfw | AGAGTACGTGGCATCCCTTG | Hernandez-Eligio et al. 2020 |
| qPCRomcCrev | CCGTTGGCATTCGTATCTCT | Hernandez-Eligio et al. 2020 |
| qPCRomcEfw | CCAGATCTGCGTGTTCTGTC | Hernandez-Eligio et al. 2020 |
| qPCRomcErev | CATGCTGCTGGACGAGTAGA | Hernandez-Eligio et al. 2020 |
| qPCRomcFfw | GACGTGGCGGCCTACATC | Rodríguez-Torres et al. 2025 |
| qPCRomcFrv | CGACGTACTCCCCGATCTTC | Rodríguez-Torres et al. 2025 |
| qPCRomcMfw | TGGAGACTACCCATGCTGAA | Hernandez-Eligio et al. 2020 |
| qPCRomcMrev | AGACGTCGAGGTGCTCGTAT | Hernandez-Eligio et al. 2020 |
| qPCRomcSfw | TCCTACCAGAACAGCAACGA | Hernandez-Eligio et al. 2020 |
| qPCRomcSrev | ATAGGAACCGCTCAGGGACT | Hernandez-Eligio et al. 2020 |
| qPCRomcZfw | AAGCCGACTGTCTCGAGTGT | Hernandez-Eligio et al. 2020 |
| qPCRomcZrev | CGGAGGTATTGATGCAGCTT | Hernandez-Eligio et al. 2020 |
| qPCRpgcAFw | GTCTCCAGAGTGCCGTAAGC | Hernandez-Eligio et al. 2022 |
| qPCRpgcARev | AGTAGGAGCCACTGCCAAGA | Hernandez-Eligio et al. 2022 |

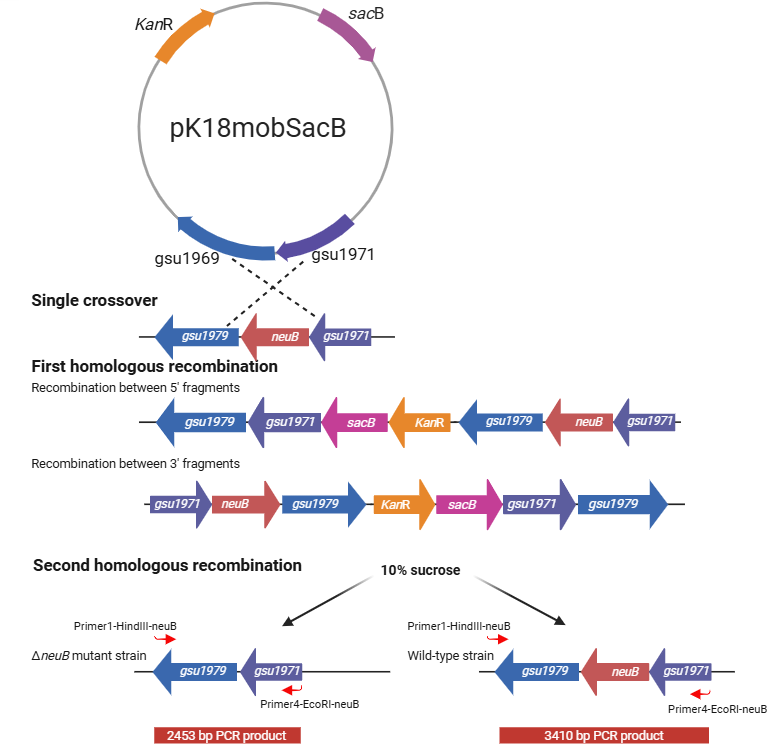

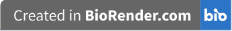

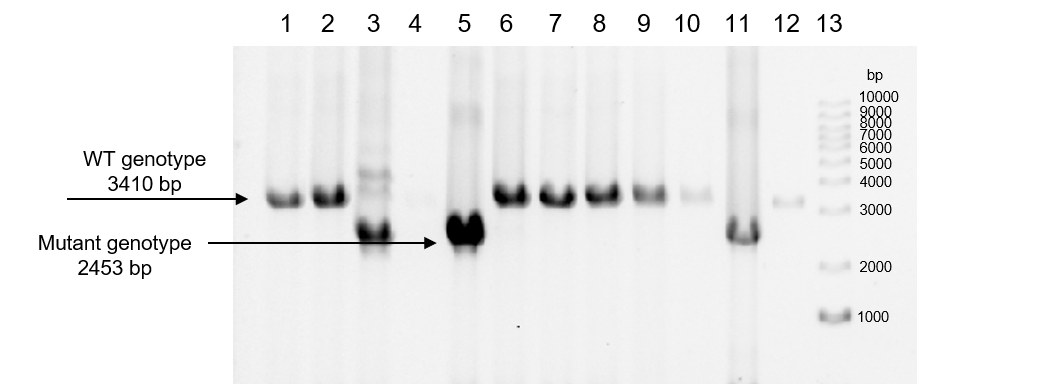

**Supplementary Fig. 1** Construction of Δ*neuB* mutant strain. Top: Diagram of the construction and verification of *neuB* gene deletion in *G. sulfurreducens*. Arrows show the primers used and expected sizes (2,453-bp for mutant and 3,410-bp for wild-type). Bottom: Agarose gel with the PCR fragments of *neuB* locus. Lanes 1–10: fragments obtained using DNA from candidate colonies of Δ*neuB* mutants; only the candidates 3 and 5 amplify the *neuB* mutant locus. Lane 11: fragment obtained using the pK18mobsacB-ΔneuB plasmid (positive control). Lane 12: fragment obtained using DNA from wild-type strain. Lane 13: 1 Kb Plus DNA ladder used as molecular weight marker (Biolabs). Arrows indicate expected molecular weights

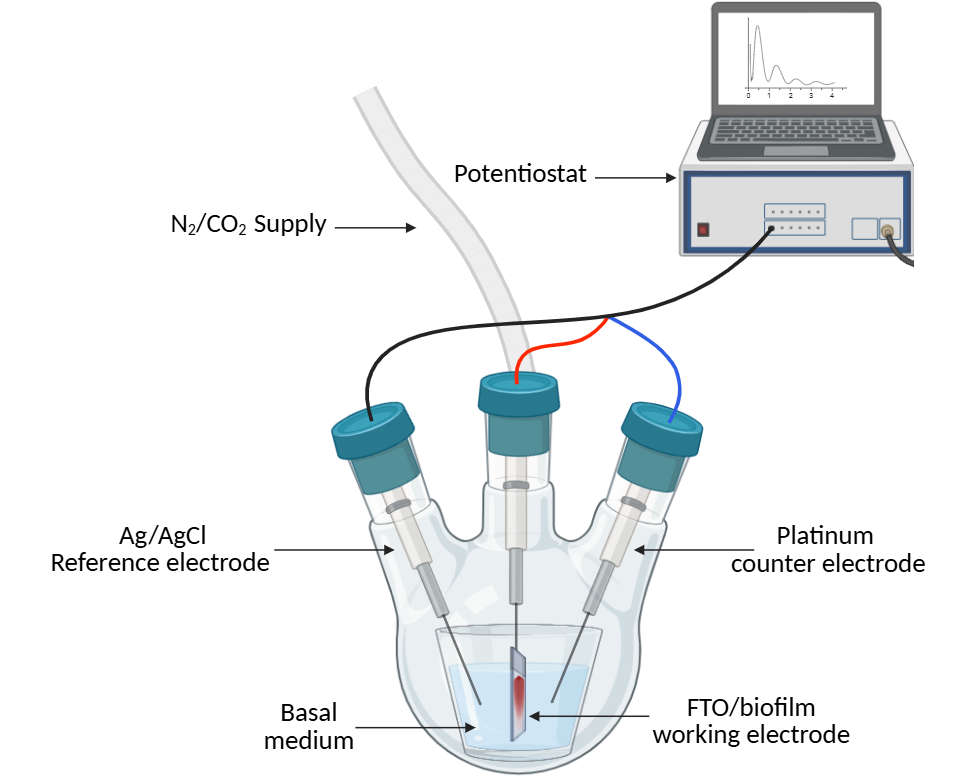

**Supplementary Fig. 2** Schematic representation of the three-electrode cell used to study the electrochemical characteristics of *G. sulfurreducens* biofilms. The setup consists of a glass cell containing a fluorine-doped tin oxide (FTO) working electrode colonized by the bacterial biofilm (center), an Ag/AgCl reference electrode (left), and a platinum counter electrode (right). The electrodes are immersed in an anaerobic basal medium and connected to a potentiostat for data acquisition (OCP, CV, and SWV measurements). Created with BioRender.com

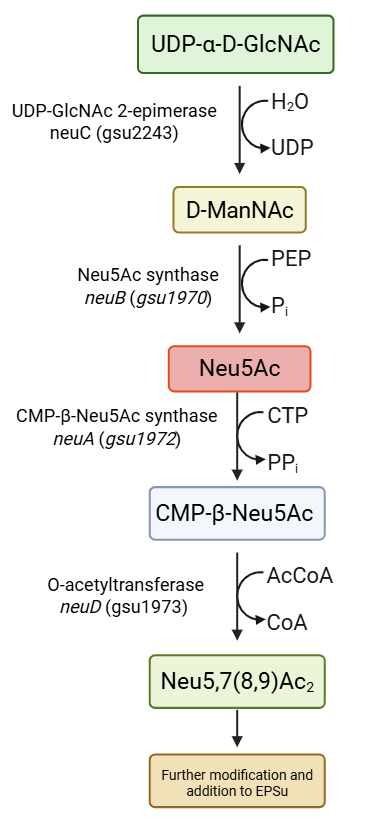

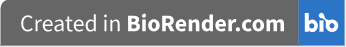

(a)

(b)

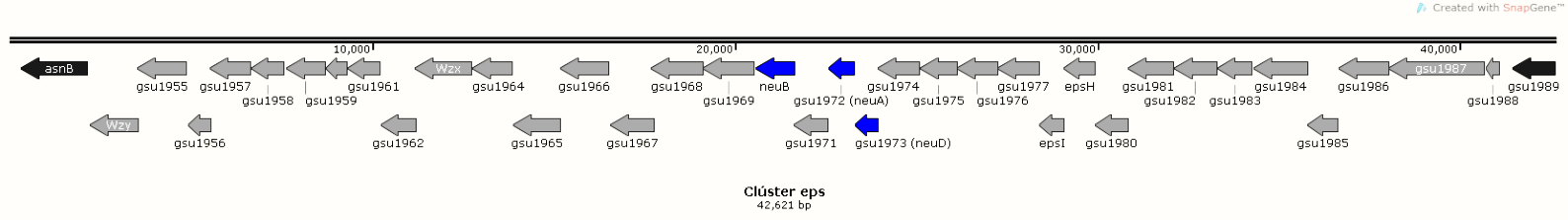

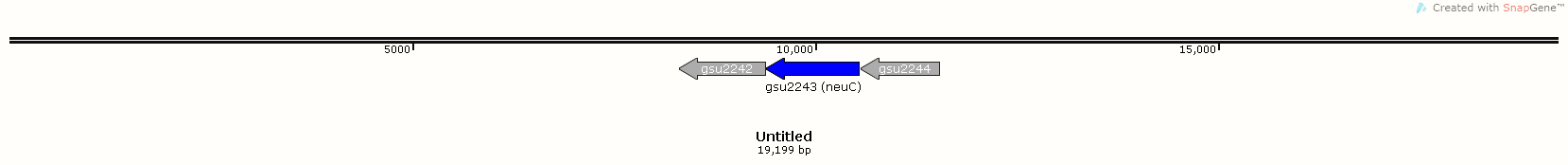

**Supplementary Fig. 3** Putative pathway of sialic acid synthesis in *G. sulfurreducens*. **a** Schematic representation of the metabolic cascade for the synthesis and modification of *N*-acetylneuraminic acid (Neu5Ac). The pathway initiates from UDP-α-D-GlcNAc and proceeds sequentially through NeuC, NeuB, and NeuA enzymatic steps to generate the activated nucleotide sugar CMP-β-Neu5Ac. Further modification by the *O*-acetyltransferase NeuD yields Neu5,7(8,9)Ac_2_ prior to its export and incorporation into the extracellular polymeric substances (EPS). The pathway scheme was adapted and modified from Ghosh (2020). **b** Genomic context and physical organization of the *eps* cluster (38,870-bp) spanning chromosomal coordinates 2,142,769-2,181,641 in *G. sulfurreducens*. Genes directly involved in sialic acid biosynthesis and modification (*neuB*/*gsu1970*, *neuA*/*gsu1972*, and *neuD*/*gsu1973*) within the *eps* cluster are highlighted in blue. The *neuC* gene (*gsu2243*), encoding the UDP-GlcNAc 2-epimerase, is located outside of this primary locus, as shown in the independent genomic region below. Arrows indicate the relative position and direction of transcription for each open reading frame. Image created with BioRender.com and SnapGene.

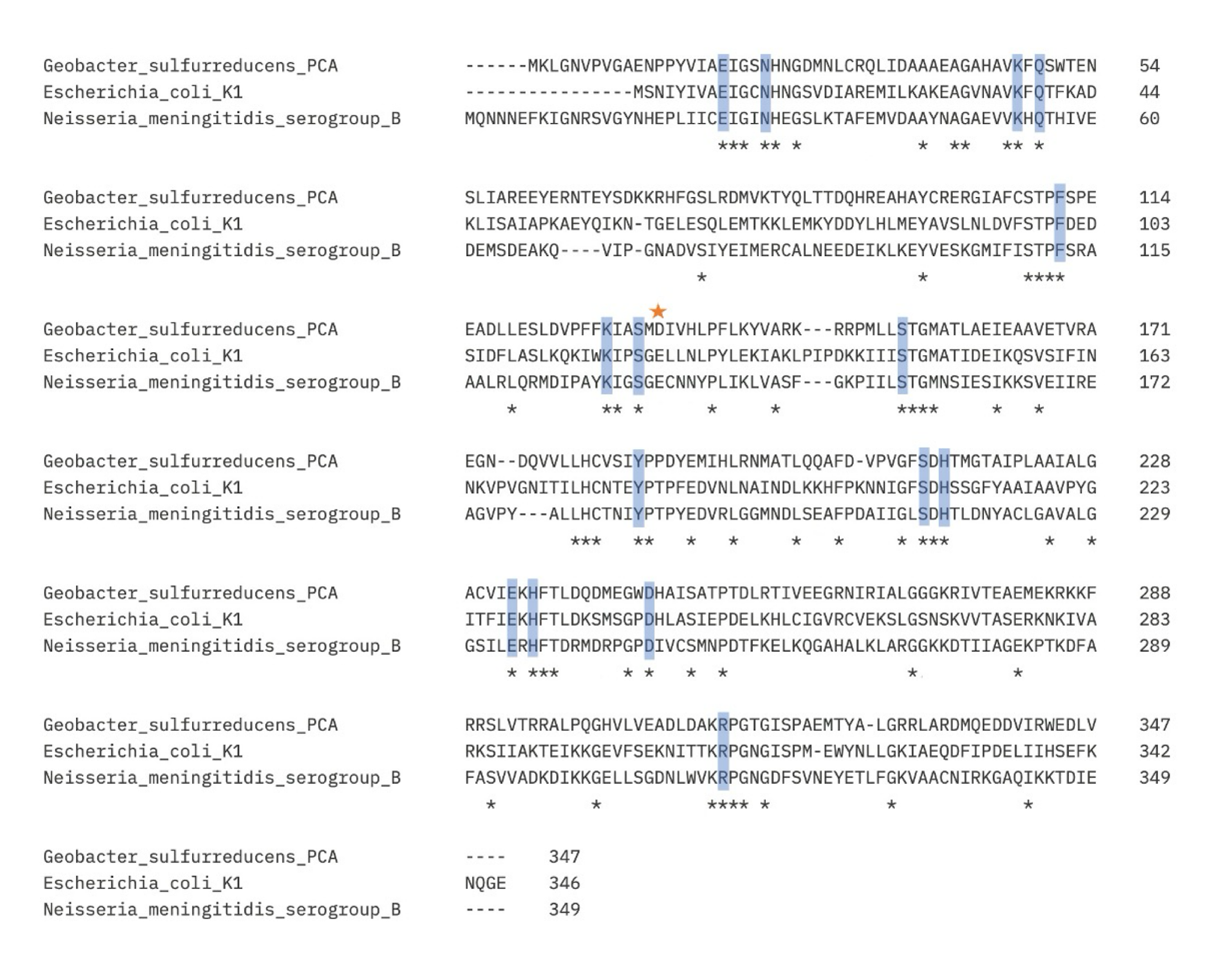

(a)

*G. sulfurreducens*

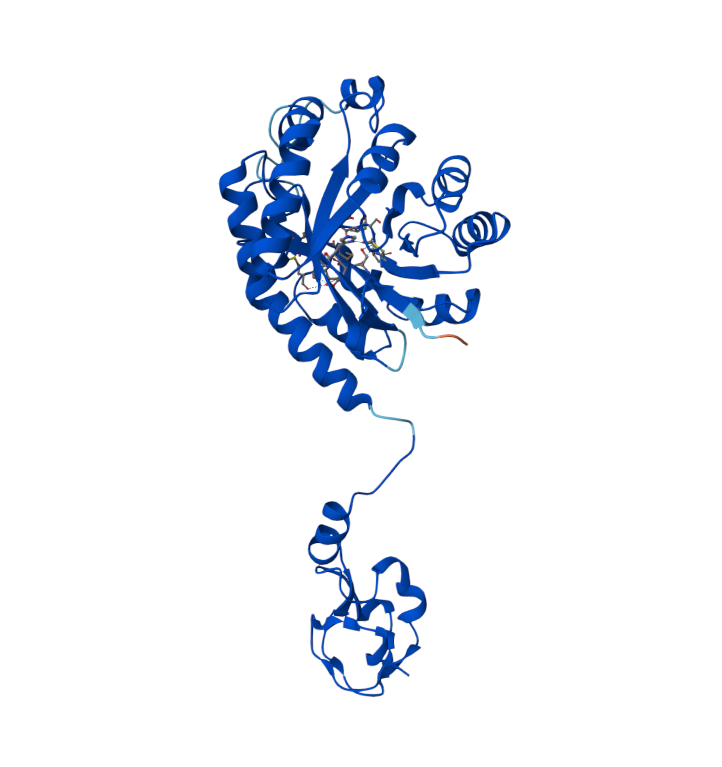

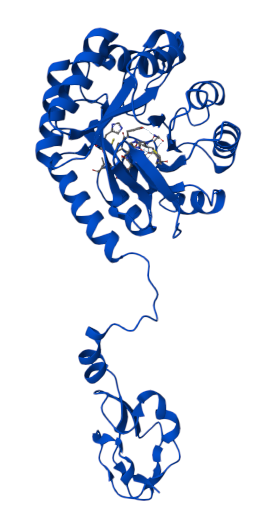

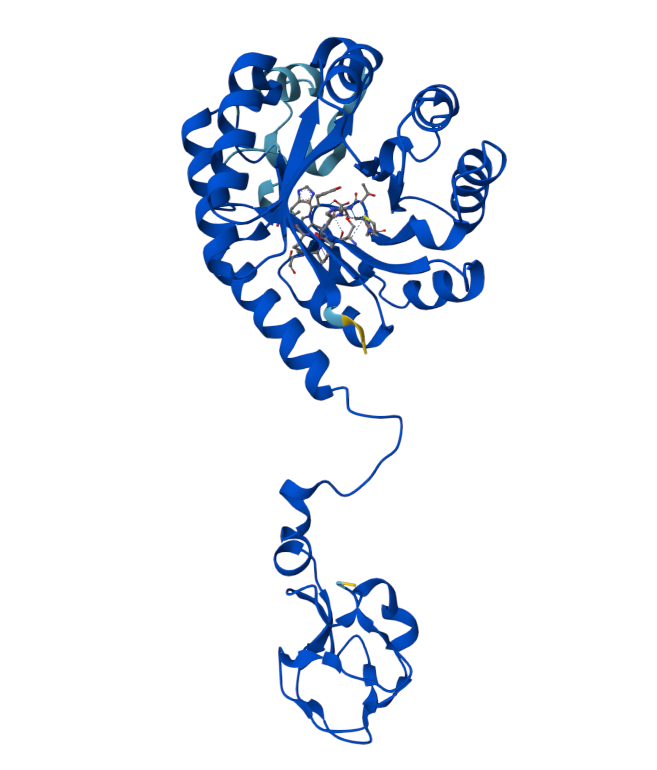

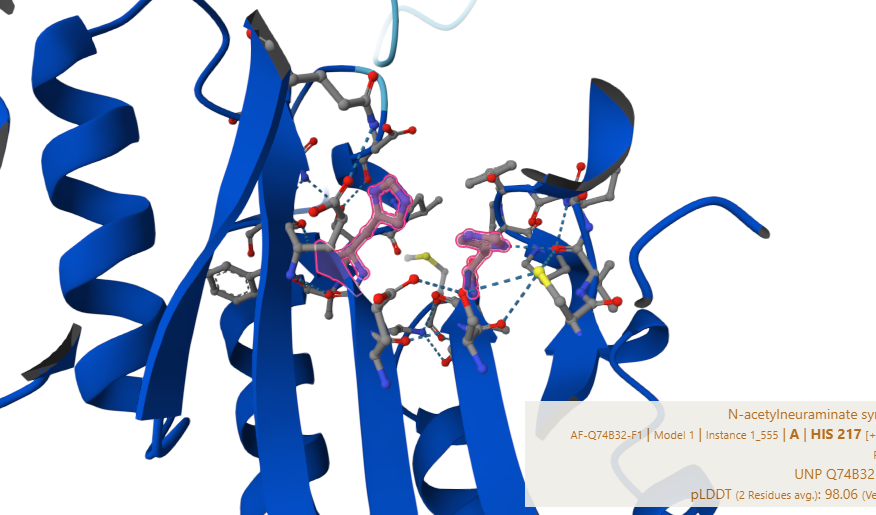

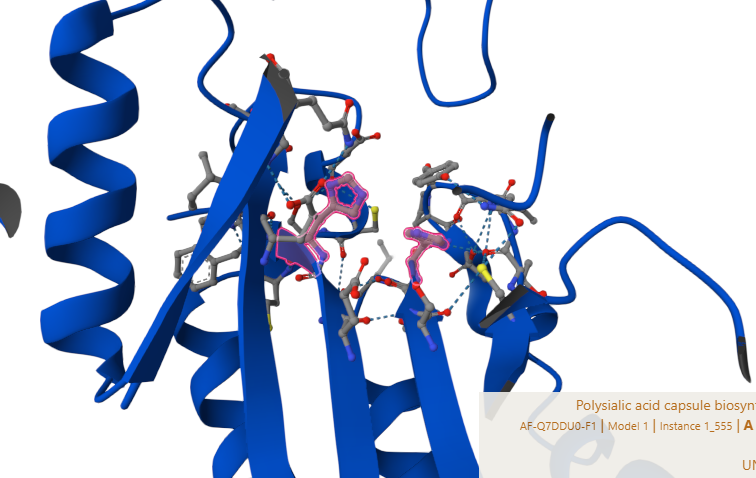

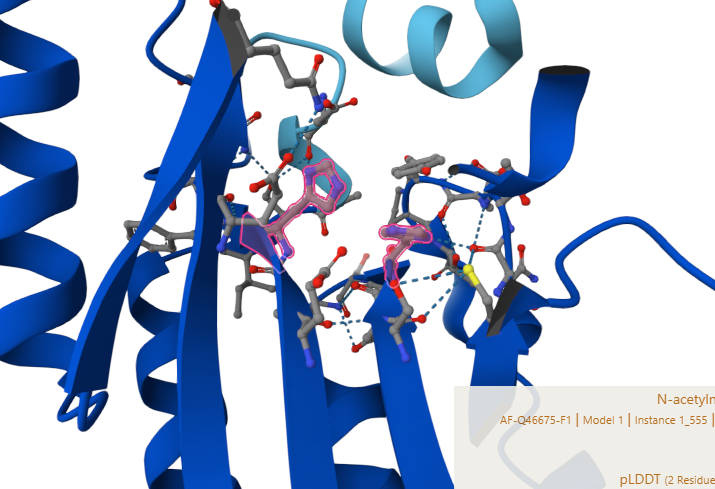

*N. meningitidis*

*E. coli*

**His 238**

**His 217**

**His 236**

**His 215**

**His 230**

**His 209**

(b)

**Supplementary Fig. 4** Analysis of NeuB protein of *Geobacter sulfurreducens.*

**a)** Multiple sequence alignment of the NeuB protein and its classical bacterial homologs. Amino acid sequence alignment comparing the N-acetylneuraminate synthase from *G. sulfurreducens* PCA (DL1) (GenBank accession: WP_235044860.1) with the sialic acid synthase from *E. coli* IHE3034 (serotype K1; GenBank accession: ADE90918.1) and the N-acetylneuraminate synthase (SynC) from *N. meningitidis* serogroup B (UniProt accession: tr|H2VFG5|H2VFG5_NEIMI). Asterisk indicate identical amino acid residues across all three sequences. Blue boxes highlight the 15 of 16 conserved amino acids involved in catalytic activity and Mn2+ metal coordination. The red star indicates the single non-conserved residue among those required for catalytic activity in *G. sulfurreducens*. Residue numbering corresponds to the *G. sulfurreducens* reference sequence. Alignment was rendered using Clustal Omega (https://www.ebi.ac.uk/jdispatcher/msa/clustalo). **b)** AlphaFold structural models of NeuB homologs illustrating the conserved dual-domain architecture—comprising an N-terminal TIM-barrel domain and a C-terminal antifreeze-like (AFL) domain—alongside overall active-site geometry. Zoomed-in insets highlight the strict structural conservation of the two essential catalytic histidine (His215 and His236) residues responsible for Mn^2+^ cofactor coordination, displaying their precise spatial orientation and respective residue numbering across the homologs, including for *N. meningitidis* (Gunawan et al. 2005).

(b)

(a)

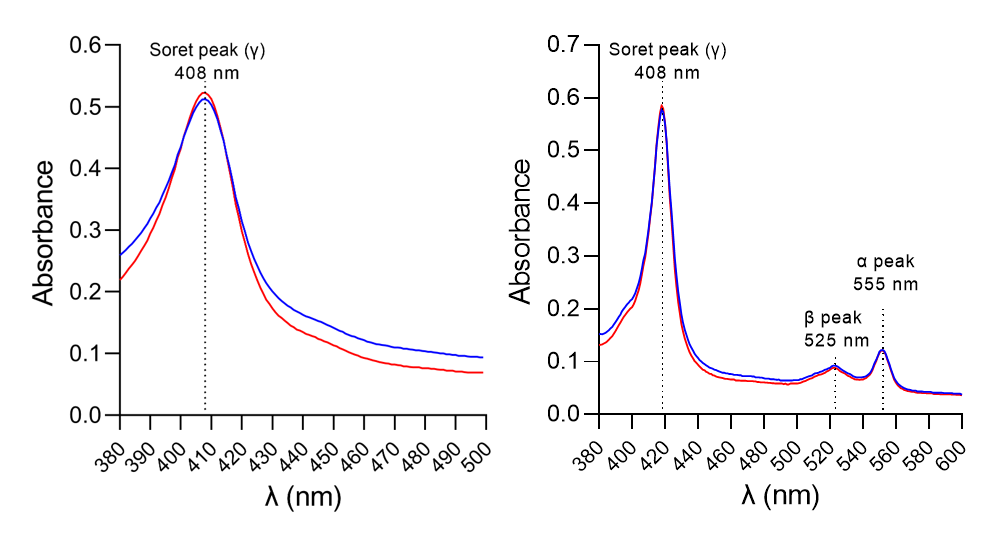

**Supplementary Fig. 5** Spectroscopic profile and relative comparison of extracellular *c*-type cytochromes. UV-vis absorption spectra of *c*-type cytochromes in the extracellular polymeric substances (EPS) of *G. sulfurreducens*. Absorbance spectra of EPS extracts from wild-type DL1 (blue line) and Δ*neuB* mutant (red line) biofilms grown on glass surfaces for 72 h. Absorbance values are normalized to cell biomass (OD_600_) to allow direct relative comparison of cytochrome c spectral profiles (Soret peak ~419 nm, β-peak ~525 nm, and α-peak ~555 nm). Data represent representative profiles from independent EPS extractions. (**a)** Spectra recorded under oxidized conditions showing the characteristic Soret (γ) band at 408 nm. (**b)** Spectra recorded under anaerobic conditions reduced with 100 mM sodium dithionite.

**References**

Caccavo F Jr, Lonergan DJ, Lovley DR, Davis M, Stolz JF, McInerney MJ (1994) *Geobacter sulfurreducens* sp. nov., a hydrogen- and acetate-oxidizing dissimilatory metal-reducing microorganism. Appl Environ Microbiol 60:3752–3759. https://doi.org/10.1128/aem.60.10.3752-3759.1994

Casadaban MJ, Cohen SN (1980) Analysis of gene control regions by gene fusion in *Escherichia coli*. J Mol Biol 138:179–207. https://doi.org/10.1016/0022-2836(80)90283-1

Chan CH, Levar CE, Zacharoff L, Badalamenti JP, Bond DR, Löffler FE (2015) Scarless genome editing and stable inducible expression vectors for *Geobacter sulfurreducens*. Appl Environ Microbiol 81:7178–7186. <https://doi.org/10.1128/AEM.01967-15>

Ghosh S (2020) Sialic acid and biology of life: An introduction. In: Elsevier eBooks, pp 1–61. doi:10.1016/b978-0-12-816126-5.00001-9

Gunawan J, Simard D, Gilbert M, Lovering AL, Wakarchuk WW, Tanner ME, Strynadka NCJ (2005) Structural and Mechanistic Analysis of Sialic Acid Synthase NeuB from *Neisseria meningitidis* in Complex with Mn2+, Phosphoenolpyruvate, and N-Acetylmannosaminitol. J Biol Chem 280(5):3555-3563. doi:10.1074/jbc.M411942200

Hernández-Eligio A, Pat-Espadas AM, Vega-Alvarado L, Huerta-Amparan M, Cervantes FJ, Juárez K (2020) Global transcriptional analysis of *Geobacter sulfurreducens* under palladium reducing conditions reveals new key cytochromes involved. Appl Microbiol Biotechnol 104:4059–4069. https://doi.org/10.1007/s00253-020-10515-0

Hernández-Eligio A, Huerta-Miranda GA, Martínez-Bahena S, Castrejón-López B, Miranda-Hernández M, Juárez K (2022) GSU1771 regulates extracellular electron transfer and electroactive biofilm formation in *Geobacter sulfurreducens*: Genetic and electrochemical characterization. Bioelectrochemistry 145:108101. https://doi.org/10.1016/j.bioelechem.2022.108101

Holmes DE, Chaudhuri SK, Nevin KP, Mehta T, Methe BA, Liu A, Ward JE, Woodard TL, Webster J, Lovley DR (2006) Microarray and genetic analysis of electron transfer to electrodes in *Geobacter sulfurreducens*. Environ Microbiol 8:1805–1815. https://doi.org/10.1111/j.1462-2920.2006.01065.x

Kim BC, Leang C, Ding YH, Glaven RH, Coppi MV, Lovley DR (2005) OmcF, a putative c-type monoheme outer membrane cytochrome required for the expression of other outer membrane cytochromes in *Geobacter sulfurreducens*. J Bacteriol 187:4505–4513. https://doi.org/10.1128/JB.187.13.4505-4513.2005

Rodríguez-Torres LM, Huerta-Miranda GA, Martínez-García AL, Mazón-Montijo DA, Hernández-Eligio A, Miranda-Hernández M, Juárez K (2025) Influence of support materials on the electroactive behavior, structure and gene expression of wild type and GSU1771-deficient mutant of *Geobacter sulfurreducens* biofilms. Environ Sci Pollut Res Int 32:16740–16759. https://doi.org/10.1007/s11356-024-33612-3

Simon R, Priefer U, Pühler A (1983) A broad host range mobilization system for in vivo genetic engineering: transposon mutagenesis in gram negative bacteria. Bio/Technology 1:784–791. https://doi.org/10.1038/nbt1183-784
